# Engineered endocrine objects tolerate hypoxia in macroencapsulation devices and enable subcutaneous engraftment in diabetic mice

**DOI:** 10.64898/2026.11.10.765512

**Authors:** Kecheng Wang, Kento Okada, Min Jin, Jerry Deng, Bo Wang, James Arthur Flanders, Phuong L. Tran, Minseok Lee, Minglin Ma

## Abstract

Islet encapsulation has been pursued for decades as a potential immunosuppression-free transplantation strategy to treat type 1 diabetes (T1D). The encapsulation device protects islets from immune cell-contact mediated killing, but it also prevents vascular penetration and intra-islet vascularization, leading to poor mass transfer and islet central necrosis. In healthy pancreas, islets with sizes > ∼40 µm are vascularized while non-islet endocrine cell clusters below ∼40 µm exist without internal capillaries. Here, we developed scalable and reproducible engineered endocrine objects (EEOs) as small as ∼47 µm by mesh-induced fragmentation of native islets, followed by a recovery culture. Under hypoxic culture condition (1% O_2_ for 24 hours), EEOs showed enhanced viability of >90%, compared to 49% for the native islet control. With a flat, thin (0.5-mm-thick) alginate-based macroencapsulation device, rat EEOs at a marginal dose of 300 islet equivalents (IEQ) achieved glycemic control for over 5 weeks in the challenging, hypoxic subcutaneous space of immunocompetent diabetic mice. The engrafted mice displayed near-physiological glycemic profiles upon glucose challenge, and the retrieved EEOs showed enhanced viability with significantly reduced central necrosis compared with native islets. Overall, engineering hypoxia-resistance into cellular products mitigates intrinsic constraints of encapsulation devices, offering a promising strategy for convenient, immunosuppression-free subcutaneous islet transplantation.

## Introduction

Type 1 diabetes (T1D) is a chronic autoimmune disorder characterized by the destruction of insulin-producing β cells and affects an estimated 9.5 million individuals globally, including 1.8 million under the age of 19 with incidence rising rapidly (1–3). Although automated insulin delivery systems have improved glycemic management, they cannot recapitulate endogenous β cell function, leaving patients with persistent risk of severe hypoglycemia and long-term complications (4, 5). Islet transplantation represents the most physiological approach to restoring β cell function. Since the Edmonton Protocol first achieved insulin independence through intraportal infusion of cadaveric islets (6, 7), the field has advanced to the 2023 FDA approval of Lantidra, the first allogeneic islet cell therapy (8). More recently, stem cell-derived islets have offered a potentially unlimited cell source, with Vertex’s zimislecel (VX-880) achieving insulin independence in most treated patients in a phase 1/2 trial (9). Despite these milestones, suboptimal survival of transplanted islets persists regardless of the cell source (10), and lifelong systemic immunosuppression remains necessary to prevent graft rejection, exposing patients to serious long-term complications and motivating strategies that protect grafts without systemic immunosuppression (11, 12).

Cell encapsulation has emerged as a promising immunosuppression-free strategy by providing a semipermeable barrier that allows the diffusion of nutrients, oxygen and insulin, while blocking direct immune cell contact (13–19). However, this immunoisolation barrier also prevents revascularization, forcing encapsulated islets to rely solely on passive oxygen diffusion from surrounding tissue. This oxygen limitation is especially problematic at the subcutaneous transplantation site, which is attractive for its clinical accessibility, safety and ease of device retrieval but is inherently hypoxic and poorly vascularized, compromising islet survival and function (20–22). Various strategies have been focused on device design to improve oxygen supply, including oxygen reservoirs and oxygen-generating materials or devices, yet these approaches remain constrained by the limited oxygen delivery capacity or the management complexity of electronic components (23–30). The prevascularization strategy, which induces blood vessel growth and increases local oxygen tension prior to transplantation, has enabled long-term subcutaneous function of macroencapsulated islets in mice but still could not prevent central necrosis in large islets (>150 µm) (31–33) and required a higher curative dose (34). This limitation is particularly relevant to the intrinsic size distributions of available β cell sources. Although small islet populations (<150 µm) exhibit higher hypoxia resistance and improved transplantation outcomes (10, 35–47), clinically approved cadaveric native islets exhibit substantial size heterogeneity, ranging from 50 to 500 µm in diameter (34, 46, 48), with large islets (>150 µm) accounting for approximately 80% of total islet volume despite representing only 20% of total islet number (38). Stem cell-derived β cell clusters, a promising alternative cell source currently under clinical investigation, are also heterogeneous when generated in suspension culture, often exceeding 150 µm and facing similar diffusion constraints (49, 50). Furthermore, necrotic islet cells can release damage-associated molecular patterns (DAMPs) and generate oxidative and inflammatory signals, potentially compromising the function of the whole graft (51–53). Collectively, achieving robust and durable survival of encapsulated islets in the subcutaneous space remains an outstanding challenge.

A newly emerged term, endocrine objects, defined as contiguous structures containing one or more endocrine cells (54–56), was introduced to encompass pancreatic endocrine structures across full size range, from single endocrine cells and small endocrine cell clusters to internally vascularized islets. Recent studies have shown that small endocrine objects that measure approximately 40 µm or less in diameter exist in the pancreas without internal vasculature (55, 57). These findings suggest an association between endocrine object size and direct vascularization, and provide a physiological rationale to engineer small endocrine objects for transplantation within encapsulation devices that are lack of vascular ingrowth. Translating this physiological architecture into an encapsulated graft requires reliable control over the size of available β cell sources. However, existing approaches to engineering islet sizes, such as enzymatic dissociation followed by reaggregation into pseudo-islets and direct islet fragmentation, remain limited by reproducibility issues, scalability and size constraints (58–60) or poorer transplantation outcomes compared with unmodified native islets (36, 61, 62).

Here, firstly, we developed a simple, yet versatile nonenzymatic method for controllable and reproducible generation of small engineered endocrine objects (EEOs) by utilizing an “islet downsizer” (Fig.1A), that mechanically refines a population of isolated pancreatic islets into small clusters within seconds independent of initial islet size. By employing meshes with different pore sizes in the islet downsizer in one step or sequentially, and followed by recovery culture, native islets with a broad size distribution were refined into small, uniform EEOs with average diameters of ∼78±4 µm and ∼47±3 µm, respectively. The recovery allows EEOs to regain glucose-responsive insulin secretion (Fig.1B) while maintaining superior viability under hypoxic conditions in vitro (Fig.1C). Secondly, we designed a geometry-optimized, flat, alginate hydrogel-based macroencapsulation device for subcutaneous transplantation of islets or EEOs (Fig. 1D). In a xenogeneic, immunocompetent diabetic mouse model, EEOs within this device restored normoglycemia for over 5 weeks in the subcutaneous space without immunosuppression. The retrieved EEO grafts showed enhanced cell survival and preserved glucose-responsive function (Fig. 1E). Together, our findings established mechanical islet downsizing coupled with recovery culture as a practical strategy for generating small, hypoxia-tolerant EEOs and revealed the importance of oxygen transport control at the intra-islet scale to mitigate diffusion constraints of encapsulation devices, offering a foundational step towards immunosuppression-free, subcutaneous β-cell transplantation for T1D.

**Fig. 1:**
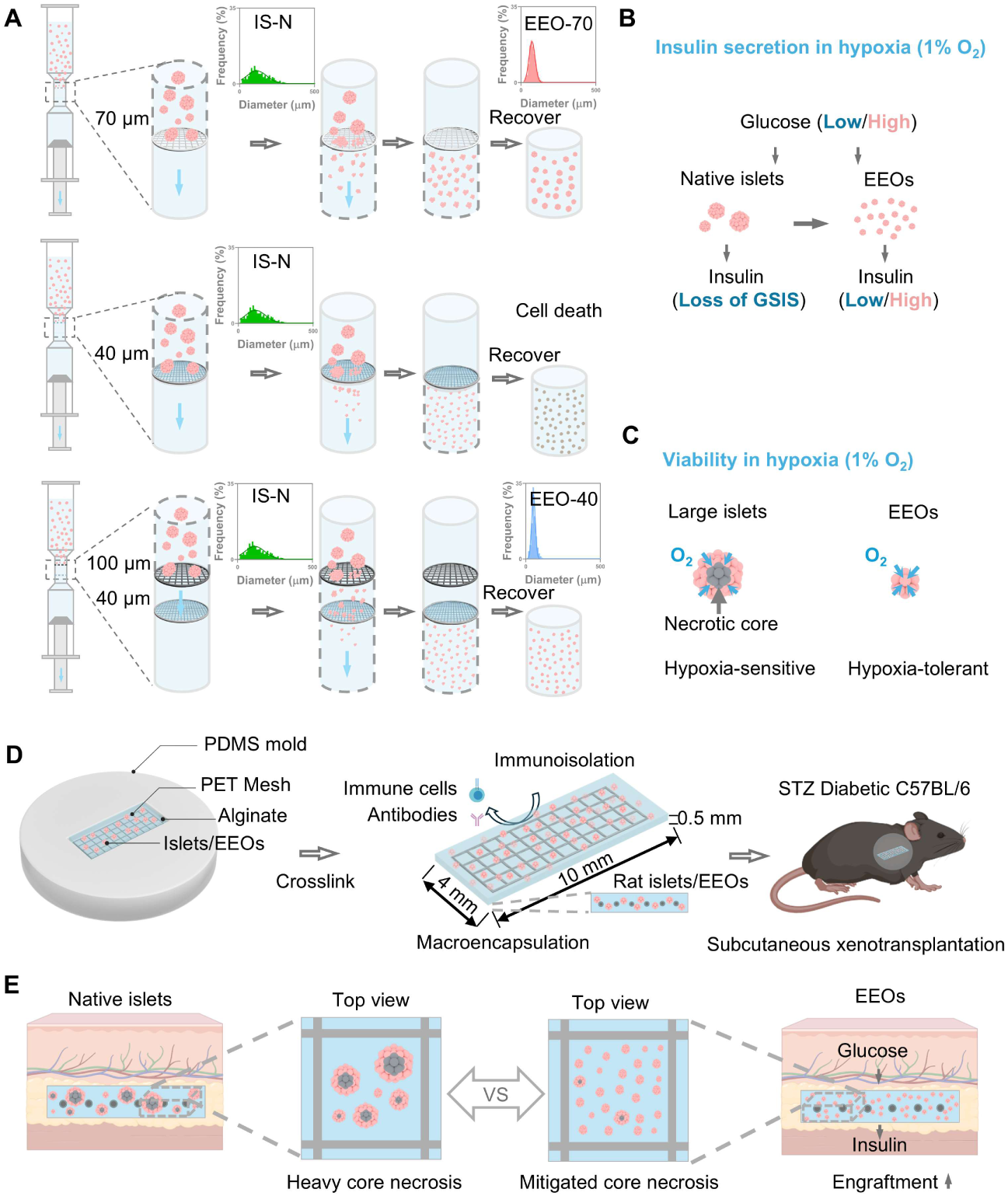
Schematics of the islet downsizing for subcutaneous engineered endocrine object transplantation. (A) Illustration of the islet downsizer, islet downsizing procedure and size distribution of native islets (IS-N) and engineered endocrine objects (EEOs), EEO-70 and EEO-40. Islets were dispersed in the medium and forced to cross the mesh with the plunger moving down. The upper, middle and bottom rows demonstrate the islet downsizing process via 70-µm, 40-µm, and dual 100-µm-40-µm nylon mesh respectively. (B) Insulin secretion study of islets and EEOs in hypoxia (1% O_2_). (C) Viability of islets and EEOs of different sizes in hypoxia (1% O_2_). (D) Illustration of islets or EEOs macroencapsulated in a flat alginate device, designed for subcutaneous xenotransplantation in STZ-induced diabetic C57BL/6J mice. (E) Illustration of native islets and EEOs in the retrieved device. The latter shows better cell survival and engraftment rate. Figures were partially created with BioRender.com.

## Results

### Computational modeling predicts enhanced hypoxia tolerance in smaller endocrine objects

The subcutaneous space is considered a relatively hypoxic environment with reported oxygen tension ranging from 1.4 to 48.5 mmHg (32, 63–70) (Table S1). This challenge is compounded in cell encapsulation devices, where the absence of revascularization leaves oxygen delivery entirely dependent on passive diffusion, driving local oxygen tension even lower. We performed computational simulations using COMSOL Multiphysics to predict the effect of endocrine object size on cell viability across peripheral oxygen tensions. For each endocrine object diameter, assuming a uniform oxygen tension at the periphery, we determined the minimum peripheral oxygen tension required to keep the endocrine object center above the survival threshold (0.08 mmHg) (34) to prevent necrotic core formation (Fig. 2A). Model parameters and boundary conditions are summarized in Table S2. Oxygen tension decreased from the endocrine object periphery toward the center, and the magnitude of this decrease strongly depends on endocrine object size. Notably, endocrine objects with diameters larger than ∼150 μm begin to develop central necrosis at a peripheral oxygen tension of 7.6 mmHg (1% O_2_) (Fig. 2B), whereas endocrine objects with a diameter of ∼50 µm remain free of necrotic cores even at 0.31 mmHg, a value lower than the oxygen tensions reported in the subcutaneous space (Fig. 2C, Table S1). Consistent with previous reports, these simulations indicate that reducing endocrine object diameter minimizes the oxygen need and the necrotic core formation even under hypoxic conditions, supporting size engineering of endocrine objects as a strategy to improve post-transplant survival.

**Fig. 2:**
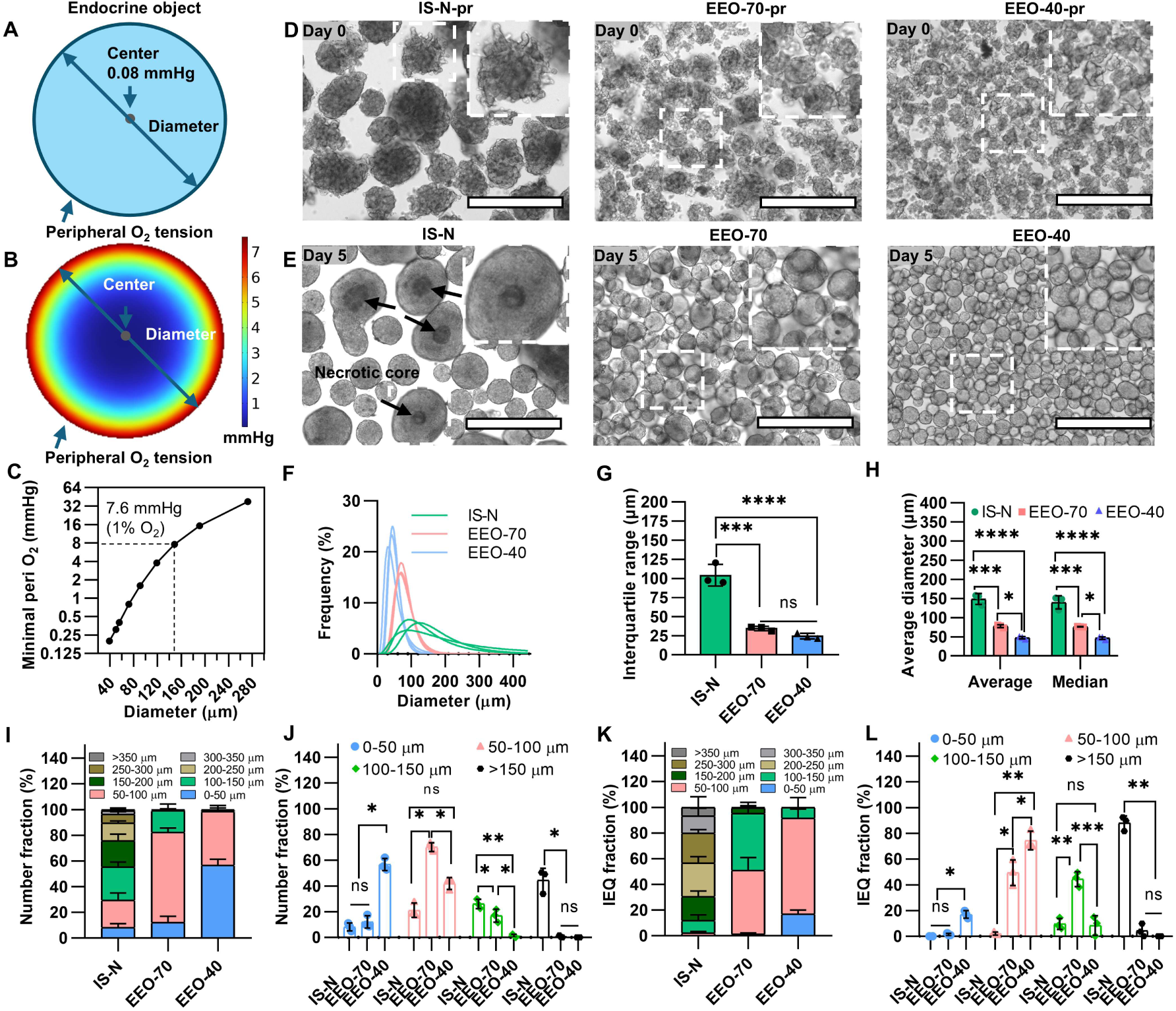
Design of the islet downsizer and characterization of engineered endocrine objects. (A) Illustration of the simulation to calculate the minimum peripheral oxygen tension to support cell survival at the endocrine object center, with a survival oxygen tension threshold of 0.08 mmHg. (B) Simulated oxygen distribution in a representative 150 µm endocrine object at a peripheral oxygen tension of 7.6 mmHg (1% O_2_) by COMSOL simulation. (C) The minimum oxygen tension required to prevent central necrosis in endocrine objects of different diameters. (D) Representative bright-field images of islets at day 0 <u>p</u>rior to recovery: native islets isolated from SD rats (IS-N-pr, left) and downsized clusters EEO-70-pr (middle) and EEO-40-pr (right). (E) The corresponding bright-field images following 5 days of recovery culture and filtration through a 20 µm cell strainer (IS-N, EEO-70, EEO-40). Scale bar, 400 µm. The arrows point to the necrotic core. (F) Diameter distribution of IS-N, EEO-70 and EEO-40, and corresponding (G) Interquartile range of size distribution (biological n=3, islet or EEO number > 290 in each group). (H) Average and median diameters of IS-N, EEO-70 and EEO-40. (I-J) Number percentage distribution and IEQ percentage distribution (K-L) of islets and EEOs across different diameter ranges. ns, not significant; *P < 0.05; **P < 0.01; ***P < 0.001; ****P < 0.0001.

### Mechanical downsizing generates size-controlled, small engineered endocrine objects with native-like morphology

The islet downsizer was designed and fabricated by mounting nylon meshes with defined pore sizes onto the Luer slip tip of a syringe (Fig. S1A). The islet downsizer allows controllable islet cell cluster production where the liquid pushes islets across the mesh powered by aspiration with the bottom plunger (Fig. S1B). To study the size controllability, nylon meshes with two pore sizes were selected: 70 µm and 40 µm. Native islets isolated from rats (Fig. S1C) were mechanically fragmented into small islet cell clusters by a single layer of 70-µm mesh within seconds (Fig. 1A, Fig. S1D). However, passing native islets through a single layer of 40-µm mesh caused islet cell cluster death (Fig. 1A). In contrast, a dual-mesh setup with a 100-µm mesh positioned above a 40-µm mesh generated small, viable islet cell clusters, with the upper mesh providing coarse pre-fragmentation (Fig. 1A). These engineered endocrine objects (EEOs) produced by 70 µm and 40 µm pore meshes collected <u>p</u>rior to recovery were named EEO-70-pr and EEO-40-pr respectively, with untreated native islets (IS-N) as the control.

Freshly isolated native islets typically exhibited rough surfaces with loosely connected cells because of enzymatic digestion and mechanical damage during the isolation process (Fig. S2A). During recovery culture, however, native islets developed a smooth and spherical morphology. Similarly, we confirmed EEO-70-pr and EEO-40-pr, though irregular and rough immediately after downsizing (day 0) (Fig. 2D), regained comparable morphology to IS-N after recovery culture of 5 days, being referred to as EEO-70-r and EEO-40-r, respectively (Fig. S2A-C). The number of IS-N decreased to ∼0.84-fold after 5 days of recovery culture, as we removed necrotized islets, whereas the number of EEO-70-r and EEO-40-r increased ∼10- and ∼18-fold, respectively (Fig. S2D-E), reflecting the fragmentation of individual native islet into multiple clusters. Importantly, the mechanical downsizing process did not compromise the yield based on islet equivalent (IEQ) calculation (Fig. S2F, Table S3). EEOs also showed the lowest batch-to-batch variation in yield, with a lower coefficient of variation (CV) than both enzymatic dissociation-based pseudo-islets (Fig. S3) and native islets (Table S4).

Following recovery culture, EEO-70-r and EEO-40-r were filtered through 20 µm cell strainers to obtain highly purified EEOs, referred as EEO-70 and EEO-40 (Fig. 2E, Fig. S2D). During the culture process, large IS-N islets developed necrotic cores, whereas EEO-70 and EEO-40 remained free of them (Fig. 2E). All groups exhibited similar surface morphology (Fig. S4A-C). EEO-70 and EEO-40 showed a narrower size distribution than native islets (Fig. 2F, Fig. S4D), reflected in a smaller interquartile diameter range (Fig. 2G) and significantly smaller average diameters (78±4 µm and 47±3 µm, respectively) than native islets (149±14 µm), along with smaller median diameters (Fig. 2H). EEO-70 and EEO-40 also showed lower inter-batch CV of diameter than native islets (Fig. S4E). Furthermore, downsizing enriched the fraction of small endocrine objects (<100 µm) in both cluster number and IEQ number. Small endocrine objects accounted for ∼29% of the total number in IS-N, rising to ∼82% in EEO-70 and ∼99% in EEO-40 (Fig. 2I-J). The corresponding IEQ fractions increased from ∼2% in IS-N to ∼51% in EEO-70 and ∼92% in EEO-40 (Fig. 2K-L). Collectively, these results demonstrate that mechanical downsizing converts heterogeneous native islets into smaller, more uniform EEOs with tunable size while preserving overall yield and native-like morphology. To explore versatility, we applied the same approach to isolated mouse islets, yielding comparable results of EEOs with preserved morphology, no central necrosis (Fig. S5A-C), and a narrow size distribution (Fig. S5D-E). These results suggest that the mechanical islet downsizing is broadly applicable across islet sources, with potential to extend to other types of cell clusters.

### Engineered endocrine objects preserve glucose-responsive insulin secretion and improve survival under hypoxia

To assess whether the downsizing process affects endocrine cellular composition and insulin secretion, we performed immunofluorescence staining, flow cytometry, and glucose-stimulated insulin secretion (GSIS) assays. EEOs overall maintained similar α, β and δ cellular composition comparable to IS-N (Fig. 3A-B, Fig. S6). For GSIS, we included pipetting-disrupted islet clusters (IS-P) as a negative control, which was previously reported to show impaired insulin secretion (61) (Fig. S7). Immediately after downsizing (day 0) prior to recovery, all mechanically processed islets, including IS-P, EEO-70-pr and EEO-40-pr, lost GSIS capacity. After 5 days of recovery, EEO-70-r and EEO-40-r restored GSIS (Fig. 3C-D). To test whether EEOs could maintain GSIS under hypoxia, we performed in vitro GSIS assays at 1% O_2_ to mimic the subcutaneous environment. Notably, GSIS was impaired in IS-N, whereas both EEO-70-r and EEO-40-r preserved adequate glucose-responsiveness (Fig. 3E-F). To explore the size effect on cell viability across oxygen conditions, we cultured IS-N, EEO-70 and EEO-40 under normoxia (21% O_2_) or hypoxia (1% O_2_) for 24 hours, followed by live/dead cell staining (Fig. 3G-H). The cell viability was characterized by the percentage by number of the healthy islets or EEOs defined as those without dead cores (Fig. 3I) and by the live area (Fig. 3J). Under normoxia, all groups showed comparable viability. Under hypoxia, however, healthy cell percentage fell sharply with size increase, from nearly 100% in EEO-40 to 80% in EEO-70 and 7% in IS-N (Fig. 3I). The live area followed the same trend (98%, 91%, and 49%, respectively) (Fig. 3J). Overall, mechanical downsizing produced EEOs that preserve cellular composition and insulin secretion with enhanced hypoxia tolerance.

**Fig. 3:**
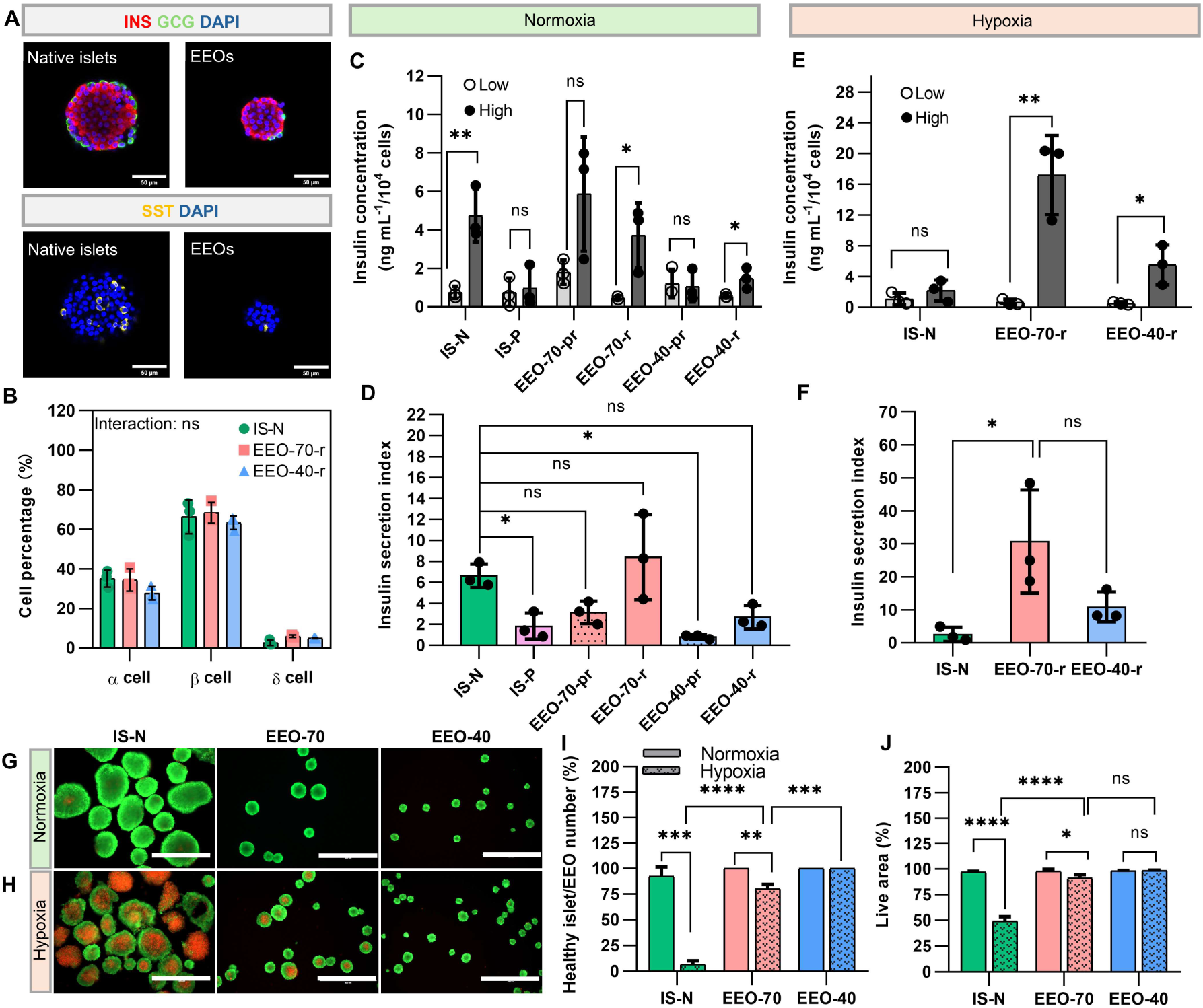
Characterization of engineered endocrine objects in cellular composition and function. (A) Representative immunofluorescence images of native and EEOs stained for INS (insulin, representing β cells, red), GCG (glucagon, representing α cells, green), SST (Somatostatin, representing δ cells, yellow) and nuclei (DAPI, blue). (Scale bar, 50 μm). (B) The percentage of α, β, δ cell numbers in native islets (IS-N), EEOs after 5 days’ recovery (EEO-70-r and EEO-40-r) by intracellular flow cytometry (biological n=3). Two-way RM ANOVA, ns (Interaction P = 0.4252). (C) In vitro glucose stimulated insulin secretion (GSIS) test of IS-N, pipetting-disrupted islet clusters on day 0 (IS-P), EEO-70-pr, EEO-70-r, EEO-40-pr and EEO-40-r in normal 21% O_2_ under low (2.8 mM) and high (16.7 mM) glucose KRB solution (biological n=3). (Abbreviations: pr, prior-to-recovery; r, post-recovery.) (D) The insulin secretion index calculated as the ratio of insulin secretion under high-glucose condition to that under the low-glucose condition. (E) GSIS of IS-N, EEO-70-r, EEO-40-r in hypoxic 1% O_2_ and (F) corresponding insulin secretion index (biological n=3). (G-H) Live/dead staining of IS-N, EEO-70 and EEO-40 (filtered EEO-70-r and EEO-40-r) under (G) 21% O_2_ and (H) 1% O_2_ (biological n=3). (Scale bar: 400 μm). Green indicates live cells and red indicates necrotic cores with dead cells. The percentage of healthy islets or EEOs (I) without a necrotic area, and the live area (J) in islets or EEOs. Data are presented as mean ± SD, analyzed by unpaired two-tailed Student’s t-test for the same group under different oxygen conditions and by one-way ANOVA with Tukey’s adjusted P value for multiple comparisons in the same oxygen condition. ns, not significant; *P < 0.05; **P < 0.01; ***P < 0.001; ****P < 0.0001.

### Encapsulated engineered endocrine objects restored glycemic control in subcutaneous xenotransplantation

To evaluate the therapeutic performance of EEOs in subcutaneous sites without immunosuppression, we macroencapsulated rat EEOs in alginate hydrogel and chose immunocompetent chemically induced diabetic C57BL/6J mice as recipients. Under hypoxic subcutaneous conditions, minimizing the diffusion distance between the surrounding tissues and the encapsulated islets is critical to maintain adequate mass transport and islet viability. Therefore, we first optimized the geometry of the macroencapsulation device. A thin, planar configuration was selected (Fig. 4A) and the device dimensions were tunable through a polydimethylsiloxane (PDMS) mold. Furthermore, a polyethylene terephthalate (PET) mesh was incorporated to provide mechanical reinforcement; the mesh geometry also provided spatial reference to identify same islets within the device before and after transplantation, allowing longitudinal monitoring of islet viability. Given that the device increases oxygen diffusion distance to encapsulated islets limiting oxygen supply, we performed computational simulations to guide the device design for in vivo studies. To assess how device thickness affects viability of encapsulated native islets and EEOs (EEO-70 and EEO-40), we applied “SHARP” model (34), a computational platform that predicts oxygen transport and cell viability in encapsulation devices by representing islets with realistic size distributions and stochastic spatial arrangements. A constant oxygen tension of 7.6 mmHg (1% O_2_) was applied at the top and bottom surfaces as the boundary condition, with an islet loading density of 15,000 IEQ/mL. Oxygen distribution was visualized in the central vertical and horizontal planes of devices with thicknesses of 0.5 mm (Fig. 4A), 0.75 mm (Fig. S8A), and 1 mm (Fig. S8B). Regions with oxygen tension below the survival threshold of 0.08 mmHg were considered necrotic. The predicted necrotic cell fraction increased with device thickness, while smaller EEOs consistently exhibited less necrosis, with necrotic fractions decreasing in the order IS-N > EEO-70 > EEO-40 (Fig. 4B). We selected 0.5 mm device thickness for subsequent in vivo studies because this thickness was predicted to keep the necrotic fraction below ∼5% in both the EEO-70 and EEO-40 groups while covering the diameter range of native islets for full encapsulation without exposure. Collectively, these simulations predict that diffusion optimization in endocrine objects and devices would mutually enhance cell viability under hypoxia, motivating subsequent in vivo validation.

**Fig. 4:**
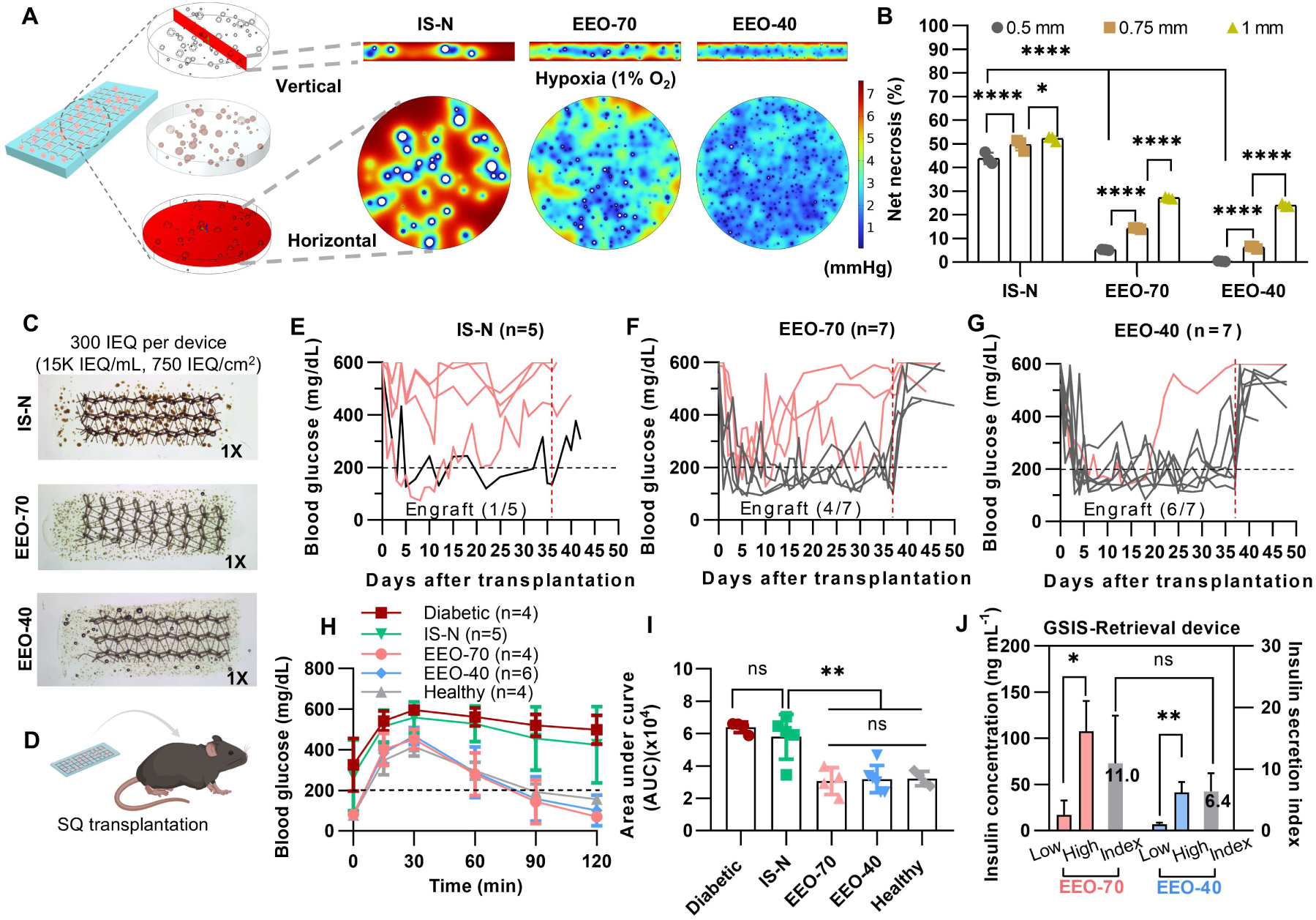
Computational modeling and in vivo evaluation of subcutaneous EEO transplantation. (A) MATLAB-generated virtual stereomicroscope images of the IS-N, EEO-70, and EEO-40 encapsulated in the 0.5-mm-thick flat alginate device and the representative oxygen tension distribution in the device indicated by the red cross-section planes simulated by COMSOL. The white area within islets indicates the necrotic area (O_2_ tension < 0.08 mmHg). (B) Predicted net necrosis rate of the islets at device thicknesses of 0.5, 0.75, and 1 mm, calculated from the white volume in (A). Data is presented as mean ± SD, analyzed by two-way ANOVA with Tukey’s adjusted P value for multiple comparisons (n = 3 iterations). (C) Representative optical photographs of device after encapsulation with a total of 300 IEQs of IS-N, EEO-70 and EEO-40 per device. (D) Illustration of subcutaneous transplantation of flat alginate devices. Blood glucose level in diabetic C57BL/6J mice after subcutaneous transplantation with (E) IS-N, (F) EEO-70, and (G) EEO-40, with engraftment rate of IS-N (1/5), EEO-70 (4/7), and EEO-40 (6/7) respectively. (H) Intraperitoneal Glucose Tolerance Test (IPGTT) of healthy mice, diabetic mice and mice with implants. Data are analyzed by two-way ANOVA with Greenhouse-Gessier’s correction and Tukey’s adjusted P value for multiple comparisons (P > 0.9999 (EEO-40 versus Healthy), P = 0.9994 (EEO-70 versus Healthy), P = 0.9994 (EEO-40 versus EEO-70), P < 0.0001 (IS-N versus Healthy), P = 0.6697 (IS-N versus Diabetic)). (I) The area under the blood glucose curve in IPGTT. (J) Insulin secretion index calculated from the ex vivo testing of retrieved functional devices of EEO-70 (n=3) and EEO-40 (n=3) in glucose stimulated insulin secretion. ns, not significant; *P < 0.05; **P < 0.01; ****P < 0.0001.

To evaluate whether the diffusion-optimized encapsulation strategy could support graft function in vivo, devices containing islets or EEOs at a dose of 300 IEQ were transplanted subcutaneously into immunocompetent diabetic C57BL/6J mice (Fig. 4C-D). A marginal dose of 300 IEQ was chosen for subcutaneous transplantation in order to provide a stringent test of graft function in the diabetic mouse model (23, 27–34, 71–73) (Table S5). Blood glucose (BG) was monitored after transplantation (Fig. 4E-G) for 5 weeks. Graft failure was defined as BG above 300 mg/dL for two consecutive days post one-week transplantation, whereas functional engraftment was indicated by BG below 200 mg/dL for two consecutive days without subsequent graft failure during the observation period. Consistent with the computational predictions, EEO groups exhibited substantially improved survival. EEO-40 group achieved the highest engraftment rate at 86% (6/7), followed by EEO-70 at 57% (4/7) and IS-N at 20% (1/5). Notably, engrafted mice in the EEO-70 and EEO-40 groups exhibited a rapid BG decline within 5 days after transplantation, suggesting early graft functionality despite the initially poorly vascularized subcutaneous environment. This rapid metabolic correction is consistent with the improved oxygen availability predicted for EEOs within geometry-optimized device. At week 5, an intraperitoneal glucose tolerance test (IPGTT) was performed to evaluate the metabolic control of blood glucose. The results revealed that all the engrafted mice in EEO-70 and EEO-40 groups were able to normalize BG levels within 120 min (Fig. 4H) and displayed near-physiological glycemic profiles comparable to healthy mice, whereas the glucose-responsiveness in the IS-N group was similar to that of non-treated diabetic mice. The area under curve (AUC) indicated almost no difference in glucose control between healthy mice and engrafted mice transplanted with EEO-70 and EEO-40 (Fig. 4I). To verify that restoration of glycemic control was graft dependent, the devices were retrieved after the functional assessment. All engrafted mice rapidly returned to hyperglycemia, confirming the in vivo function of the transplants (Fig. 4E-G). In ex vivo GSIS, EEO-70 and EEO-40 in retrieved devices showed glucose-responsive insulin secretion with comparable insulin secretion index (Fig. 4J). Overall, these results demonstrated that combining EEOs with the thin, diffusion-optimized alginate device substantially reversed hyperglycemia for at least 5 weeks after subcutaneous xenotransplantation in diabetic mice without immunosuppression, with near-physiological glucose clearance and no delayed response in IPGTT.

### Engineered endocrine objects exhibited enhanced viability after subcutaneous transplantation

To determine whether sustained function reflected improved cell survival, we retrieved all devices at week 5 and evaluated endocrine viability (Fig. S9A). Retrieved devices were all covered by a layer of connective tissue (Fig. S9B) with some visible blood vessels on the surface (Fig. S9C). The β cells were stained red by dithizone (DTZ) in EEO-70 and EEO-40 groups, identifying the islets in the retrieved device after the 5-week transplantation (Fig. S9D-F).

To directly assess changes in islet morphology and viability during transplantation, we compared brightfield images of the same device regions captured before transplantation (Tx) and after retrieval, as identified by the PET mesh pattern (Fig. 5A-B), and further analyzed retrieved islets by live/dead staining (Fig. 5C), H&E staining (Fig. 5D), and immunofluorescence staining for insulin and glucagon (Fig. 5E). Based on the brightfield images, all pre-transplanted islets or EEOs were intact and compact with smooth surfaces, which we defined as normal morphology (Fig. 5B). After transplantation, large islets from IS-N (>150 µm) showed rough surfaces, necrosis, and shrinkage with surrounding dissociated cells, whereas only small islets from IS-N retained normal morphology. In the EEO-70 and EEO-40 groups, EEOs retained their pre-transplantation morphology. We then quantified the percentage of healthy islets or EEOs, defined as those without a dark necrotic center (Fig. S10). This percentage rose markedly from 24±15% in IS-N (n = 5) to 78±17% in EEO-70 (n = 7) and 74±16% in EEO-40 (n = 7) (Fig. 5F). Necrotic areas did not differ significantly between EEO-70 and EEO-40 (Fig. 5G). Live/dead staining showed extensive cell death in large islets from IS-N but preserved viability in EEO-70 and EEO-40 (Fig. 5C). H&E staining demonstrated loss of structural integrity and predominant necrotic regions in large IS-N islets. Necrosis was rarely observed in EEO-70 islets whereas EEO-40 retained intact structure (Fig. 5D). Immunofluorescence for insulin and glucagon confirmed that EEO-70 and EEO-40 islets remained functional after retrieval (Fig. 5E). Furthermore, α-SMA and CD 31-positive vascular structures were observed in the connective tissue at the device-host interface (Fig. S11). Although these vessels did not directly vascularize islets, their proximity to device surface may support insulin transport from the graft to the host circulation, consistent with the rapid glucose response observed in IPGTT (Fig. 4H-I). Overall, these results demonstrate that EEOs tolerate hypoxia in macroencapsulation device and enabled sustained graft function in subcutaneous space.

**Fig. 5:**
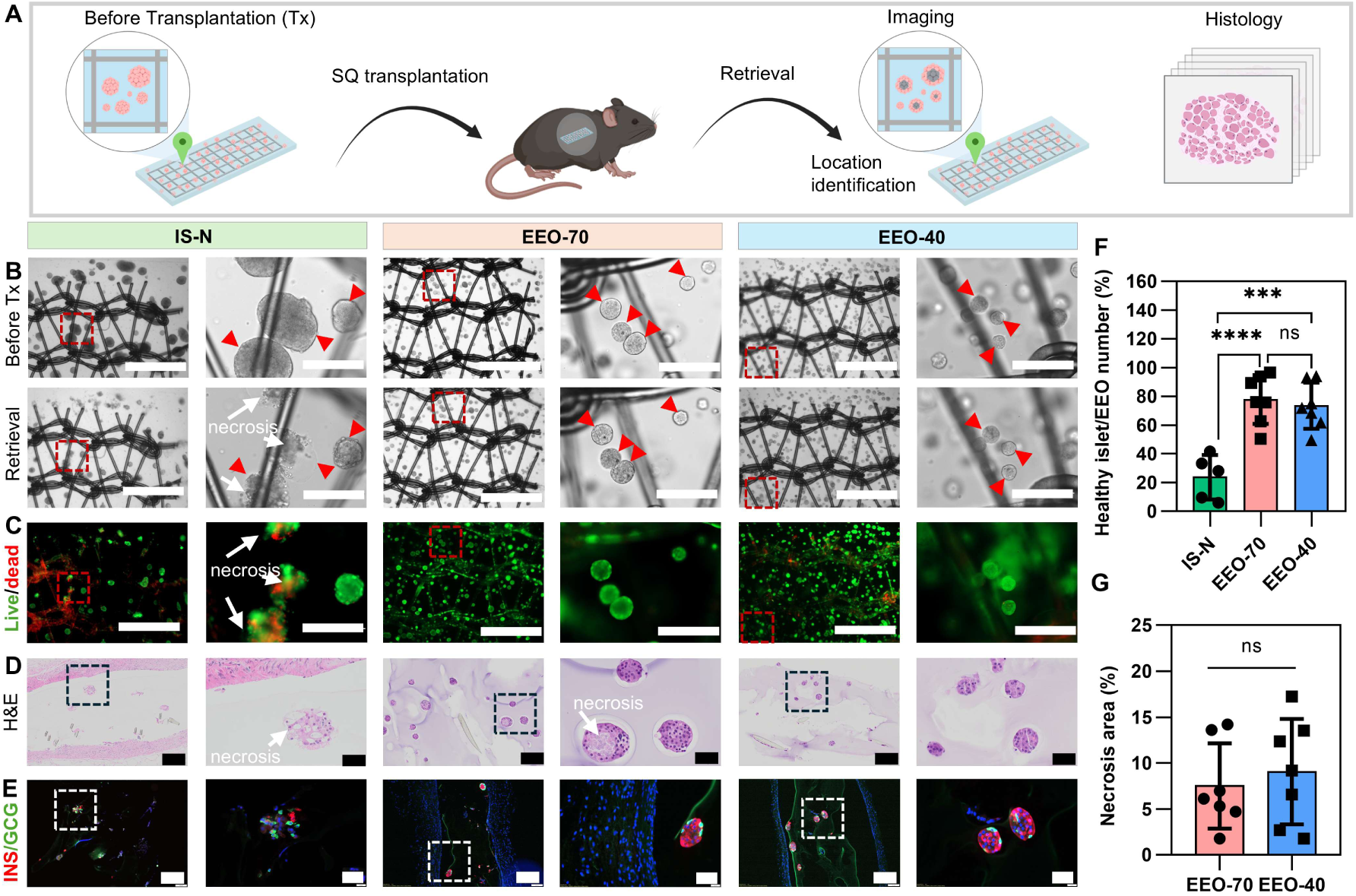
Viability and structure integrity of retrieved IS-N, EEO-70 and EEO-40. (A) Illustration of the workflow for evaluating islet graft before and after transplantation. (B) Representative bright-field images of IS-N, EEO-70 and EEO-40 in the device before transplantation (Tx) (top) and after retrieval (bottom) (Scale bar: 1000 μm) along with magnified images in the red frame (Scale bar: 200 μm). Red arrows indicate the same islets in the same location and white arrows indicate the necrosis. (C) Corresponding live/dead staining of IS-N, EEO-70 and EEO-40 in retrieved devices. (Scale bar: 1000 μm) along with magnified images in the red frame (Scale bar: 200 μm). (D) Representative H&E staining of islets in retrieved devices (Scale bar: 200 μm) along with magnified images in the black frame (Scale bar: 50 μm). (E) Immunofluorescence staining of insulin (red), glucagon (green), and nuclei (blue) in islets within retrieved devices. (Scale bar: 200 μm) along with magnified images in the white frame (Scale bar: 50 μm). (F) Percentage of healthy islets or EEOs by number in retrieved devices. IS-N (n=5), EEO-70 (n=7), EEO-40 (n=7). (G) Percentage of the EEO necrotic area of retrieved EEO-70 (n=7) and EEO-40 (n=7) devices. Data is presented as mean ± SD, analyzed by unpaired two-tailed Student’s t-test. ns, not significant; ***P < 0.001; ****P < 0.0001.

## Discussion

The facts that native islets are internally vascularized and that smaller endocrine objects (∼40 µm) without internal capillaries exist in human pancreas provide a physiological rationale for engineering small-size endocrine objects for transplantation within encapsulation devices that are lack of vascular ingrowth. We developed a mechanical downsizing approach that reduced the mean diameter of native rat islet populations from ∼149 µm to ∼78 µm (EEO-70) or ∼47 µm (EEO-40). These engineered endocrine objects maintained high viability and preserved GSIS under hypoxia (1% O₂). When encapsulated in a flat, 0.5-mm-thick alginate devices, they restored normoglycemia following subcutaneous transplantation in immunocompetent diabetic mice and exhibited significantly less central necrosis than native islets at graft retrieval 5 weeks after transplantation.

Subcutaneous transplantation of encapsulated islets remains challenging because the site is poorly vascularized causing hypoxia and the encapsulation devices exclude vascular penetration imposing additional diffusion barrier. Although subcutaneous prevascularization could increase local oxygen tension, it could not prevent the proportion of large islets (>150 µm) from developing central necrosis in the encapsulation device (32). Islet size engineering provides a strategy to improve islet hypoxia tolerance. However, a reproducible method to engineer heterogenous sized islets to small sizes remained challenging. The most widely used method for islet size engineering involves enzymatic dissociation of islets into single cells, followed by reaggregation to form uniformly sized pseudo-islets (44, 74–76) (Table S6). However, enzymatic islet dissociation remains difficult to standardize and scale because islet size varies widely within preparations and among donors. Since dissociation progresses from the periphery toward the core, this heterogeneity results in size-dependent differences in dissociation kinetics. Moreover, the narrow window for achieving sufficient dissociation without compromising cell viability, together with the absence of an objective endpoint, limits process control and the reproducibility of pseudo-islet production (58, 59). Direct islet fragmentation generated small islet clusters; however, these clusters yielded poorer transplantation outcomes than unmodified native islets (36, 61). In the present study, we demonstrated the proof-of-concept of enzyme-free, mechanical approach for islet size engineering, which robustly processes the whole population of islets via a single mechanical pass through a mesh within seconds by fluid pressure. The “islet downsizer” has been built based on a pair of syringes, which provides operation control. Although previous studies reported impaired islet function following mechanical fragmentation (36, 61), we identified that the impairment in our system was transient and reversible, with recovery culture enabling the restoration of GSIS. Furthermore, our mesh fragmentation method enabled the nonenzymatic generation of EEOs at the 40-µm scale, representing, to our knowledge, the smallest size reported to date using a nonenzymatic method (Table S6). Another key advantage of mechanical downsizing is the improved functional endocrine mass yield, defined here as the proportion of initially isolated islet mass that remains functional and suitable for transplantation. In native islet preparations, central necrosis in large islets during pretransplant culture reduces this yield. Mechanical downsizing converts these vulnerable islets into smaller EEOs with greater hypoxia tolerance, thereby preserving functional endocrine mass for transplantation. This approach can be readily integrated into standard islet isolation workflows to improve islet quality and support a broad range of islet studies under oxygen-limited conditions.

With our functional and hypoxia-tolerant EEOs, we explored the effect of size reduction on the survival and function of EEOs in encapsulation devices after subcutaneous transplantation. Wang et al. highlighted the oxygen limitation in subcutaneous sites, showing that macroencapsulated islets restored glycemic control in prevascularized subcutaneous sites but failed in unmodified sites in immunocompetent diabetic mice using ∼500 IEQ of rat islets at a loading density of ∼14,300 IEQ/mL (32). Consistent with their results, our study showed 4 of 5 immunocompetent diabetic mice receiving devices containing native islets at an unmodified subcutaneous site failed to achieve normoglycemia using a lower dose of ∼300 IEQ of rat islets at a similar loading density ∼15,000 IEQ/mL. In contrast, devices containing EEO-40 with a mean diameter of ∼47 µm at ∼300 IEQ dose restored and maintained normoglycemia in 6 of 7 mice for at least 5 weeks. At graft retrieval, fewer than 30% of EEOs exhibited necrotic cores, compared with more than 76% of native islets. Our findings support endocrine object size reduction enhances encapsulated graft survival and function in oxygen limited subcutaneous environment without prevascularization. Jeon et al. also reported glycemic correction without prevascularization in immunodeficient mice using ∼250 IEQ of rat islets at a low loading density of 2,500 IEQ/mL; however, transplantation of ∼500 IEQ into immunocompetent C57BL/6 mice failed to restore glycemic control without co-delivery of the antifibrotic drug GW2580 (29). This suggests that monocyte and macrophage recruitment, and pericapsular fibrosis on device surface may compromise encapsulated islet function. In contrast, at 60% of their dose and at six times their loading density, our alginate devices containing EEO-70 or EEO-40 achieved glycemic correction in diabetic C57BL/6 mice without antifibrotic drugs. Despite fibrotic capsule formation, IPGTT responses one month after transplantation closely resembled those of healthy mice without any delays. The rapid glycemic responses may also partly reflect adequate glucose, oxygen and insulin transport across the device and within EEOs. These findings suggest EEOs with a mean diameter of ∼78 µm or less can survive and function within encapsulation devices under subcutaneous hypoxia even in the presence of fibrotic capsule. Together, our work highlights engineering hypoxia-resistance into cellular products mitigates intrinsic constraints of encapsulation devices which increase diffusion barrier and prevent direct vascular access.

Successful subcutaneous transplantation of macroencapsulated EEOs requires coordinated control of oxygen transport at both the intra-EEO and device scales. Mechanical downsizing divides native isolated islets into smaller EEOs, increasing the total surface area available for oxygen diffusion and shortening diffusion distances to the core. Our simulations also indicate that smaller endocrine objects require lower peripheral oxygen tension for survival. Together, these advantages enhance the hypoxia tolerance of EEO populations. Device thickness is another critical determinant of oxygen and nutrient availability for EEOs (34, 77–79). Simulated necrosis increased with device thickness in both native islets and EEO populations, indicating that islet downsizing cannot fully overcome device-scale diffusion limitations. Central vertical cross-sections of the devices showed a common pattern of steep oxygen gradients from the device surface toward the core, consistent with previous findings (29). In contrast, central horizontal cross-sections revealed distinct oxygen distributions in islet and EEO populations. Native islets exhibited substantial central necrosis despite relatively high peripheral oxygen tension, indicating that local oxygen availability remained insufficient to sustain the survival of their cores. At the same IEQ dose, devices with smaller EEOs, particularly EEO-40, exhibited lower oxygen tension at their peripheral but more spatially uniform oxygen tension distribution. In 0.5-mm-thick devices, 1% oxygen level was sufficient to support the survival of EEO-40 populations. At a fixed cell loading density, decreasing device thickness shortens the diffusion path and improves oxygen penetration into the device interior, resulting in higher oxygen tension in the device core, while reducing the size of endocrine objects lowers the peripheral oxygen tension required for their survival. These findings support a dual-scale mechanism in which device thickness governs oxygen delivery to encapsulated EEOs, while EEO diameter determines whether the available peripheral oxygen can sustain the survival of their cores. Given the marginal dose of 300 IEQ (Table S5), the relatively high loading density of 15,000 IEQ/mL and the presence of fibrotic capsule imposing an additional diffusion barrier, improved oxygen transport is a key feature of our platform that supports functional engraftment of macroencapsulated EEOs in immunocompetent diabetic mice at unmodified subcutaneous sites. Together, these findings highlight a key design principle for EEO macroencapsulation: matching device-scale oxygen delivery with the size-dependent peripheral oxygen requirements of EEOs to sustain graft function under hypoxic conditions.

Beyond EEOs’ hypoxia tolerance and functional advances, they could also serve as standardized β-cell preparations to reduce discrepancies across β-cell transplantation studies. Pancreatic islets lack a universally accepted size definition (55), and their size varies widely within preparations and among donors (59). This variability results in each isolated islet preparation having a different size distribution, leading to variable transplantation outcomes and complicating comparisons across studies. With our approach, islet isolated from pancreas could be converted to functional EEOs with a narrow, reproducible size distribution determined by the mesh pore size. This provides a convenient way to standardize islet preparations through mesh treatment rather than setting a criterion for islet size range. Furthermore, EEOs mitigate variability from intrinsic islet size heterogeneity within preparations and among donors. Standardizing islet preparations into EEOs before transplantation could improve reproducibility, enable more reliable evaluation of β-cell replacement therapies and facilitate comparisons of transplantation outcomes across experiments and research groups.

Several aspects of the platform remain open to further optimization. Characterizing the disruption of intercellular junctions and extracellular matrix during mechanical downsizing and their restoration during recovery culture could guide protocol refinement. Comparisons between EEO-70 and EEO-40 revealed benefits and potential trade-offs of size reduction. Both outperformed native islets in hypoxia tolerance and in vivo function, with smaller EEOs showing greater survival. EEO-70 retained a subpopulation of larger clusters that developed central necrosis under hypoxia in vitro, whereas EEO-40 showed no necrotic cores. Consistently, functional engraftment rates were 6/7 for EEO-40 and 4/7 for EEO-70. However, EEO-40 exhibited lower insulin secretion during GSIS. One possible explanation is that forcing islets through smaller mesh openings exposes a greater proportion of cells to mechanical stress. The current islet downsizer employs commercially available nylon mesh with relatively thick fibers (∼35 µm of fiber width in 70 µm pore mesh and ∼20 µm in 40 µm pore mesh). Custom meshes with thinner fibers may reduce mechanical injury and improve the balance between islet survival and function. We have demonstrated the versatility of mechanical islet downsizing using rat and mouse islets. Further validation will involve human cadaveric islets and stem cell-derived islets. For clinical-scale processing, the mesh area could be adjusted to accommodate different doses (Fig. S12). At the current processing density of 44,000 IEQ per cm² of mesh area, processing 500,000 IEQ would require approximately 11 cm² of mesh (3.8 cm in diameter) in a scaled-up downsizer. Integrating established 3D culture platforms, including suspension bioreactors and specialized media with additives may support large-volume recovery culture while minimizing unwanted cluster fusion (80–82). Beyond thickness, device optimization could include increasing the surface-to-volume ratio to facilitate intra-device oxygen transport (28). Future in vivo studies could incorporate prevascularization (32), antifibrotic modifications (83, 84) or immunomodulatory cells (85) to modulate the host fibrotic responses and support long-term graft function.

In summary, mechanical downsizing offers a simple, robust approach to generate small, functional EEOs with enhanced resistance to hypoxia-induced necrosis and a convenient way to standardize islet preparations for transplantation. Integrated with a thin, planar alginate macroencapsulation device, EEOs mitigated intrinsic constraints of encapsulation devices and enabled sustained glycemic control following subcutaneous transplantation in immunocompetent diabetic mice without prevascularization, external oxygen supplementation, or antifibrotic drugs. These findings highlight a key design principle of matching device-scale oxygen delivery to the size-dependent oxygen requirements of encapsulated EEOs. This platform provides a practical basis for further development of subcutaneous, immunosuppression-free β-cell transplantation for T1D treatment.

## Materials and Methods

### Animals

C57BL/6J mice (6-12-week, male, 25-30g) were purchased from The Jackson Laboratory. Sprague-Dawley (SD) rats (7-8-week, male or female, ∼250g) were purchased from Charles River Laboratories. All animal procedures were approved by Institutional Animal Care and Use Committee (IACUC) at Cornell University (protocol number: 2023-0086).

### Fabrication of islet downsizer

Nylon meshes were obtained from 100 μm, 70 μm and 40 μm cell strainers (VWR®, 76327). The meshes were mounted on the outlet of syringes by an ultrasonic plastic spot welder followed by ethylene oxide sterilization for use.

### Preparation and purification of engineered endocrine objects (EEOs)

Islet isolation process was presented in Supplementary methods. For EEO-70 preparation, a 70 μm nylon mesh was pre-welded to a 3 mL syringe outlet, which connected with the other 3 mL empty syringe via a luer lock connecter as shown in Fig. 1A. First, the syringes and nylon mesh were thoroughly rinsed with 1mL of complete RPMI medium (with 10% FBS and 1% penicillin/streptomycin) by pulling the lower syringe plunger, leaving 0.5 mL of medium in the upper syringe. Then, the islets suspension was loaded into the upper syringe until the total volume reached 2 mL. Next, the piston of the lower syringe was pushed upward to dislodge the islets that settled at the bottom of the upper syringe. A pipette (fitted with a 200 µL tip) was used to disperse all the islets, and the bottom syringe plunger was pulled down to force the culture medium with islets passed through the mesh within one second (at a flow rate of ∼2 mL/s). Subsequently, all islets were downsized to islet clusters with uniform size and then transferred into a 10 cm petri dish. The conjoined syringe device was rinsed with an additional 2 mL of culture medium in the upper syringe and repeated the above steps to collect all the residual islets in the syringes. Each dish contained about 500 IEQ EEOs and was cultured in the incubator at 37 °C (day 0). Purification was conducted on the next day (day 1). Firstly, EEOs were collected and centrifuged at 230G for 2 min in a 15mL centrifuge tube (tube1). 1.5 mL of

Histopaque-1077 was added to a new 15-mL centrifuge tube (tube2), then 1mL of islet suspension from tube1 was gently transferred to the top of the Histopaque solution. Tube1 was rinsed with 1mL of complete RPMI to collect all the remaining residual EEOs which were loaded on top of tube2. Gradient density centrifugation at 400G was performed for 5 min at 1 acceleration and 0 deceleration. The top layer supernatant of 1 mL was removed. All the EEOs at the interface between the RPMI medium and the Histopaque-1077 were collected and added to 5 mL washing buffer (RPMI with 2.5% FBS and 1% penicillin/streptomycin), followed by centrifugation at 230G for 2min. Surprenant was aspired and 5 mL washing buffer was added to suspend islets, followed by centrifugation at 230G for 2min. This procedure was repeated one more time. The purified EEOs were then transferred to a 10 cm petri dish with 10 mL complete RPMI medium, and the center of the petri dish was gently pipetted four times to prevent EEO aggregation during the incubation. On day 3, the same purification process was repeated to remove dead cells in the medium.

For EEO-40 preparation, the upper syringe’s outlet was mounted with 100 μm nylon mesh, and a luer lock connector in the middle and the lower syringe was mounted with 40 μm nylon mesh. The larger mesh was applied to refine the islet size suitable for preparing smaller islet clusters, since directly passing large islets through the 40 μm pores caused significant cell death. A stepwise downsizing minimized the damage on islets. The remaining procedures and purification methods were the same as those islets downsized using the 70 μm mesh.

### Characterization of EEOs

On day 5, EEOs were characterized by IEQ yield and static glucose stimulated insulin secretion (GSIS) (86) (Supplementary method). In vitro GSIS assays were performed using rat islets (biological n = 3). Islets isolated from each donor were split into three portions, with one retained as the native control and the other two used to prepare EEO-70-r and EEO-40-r, respectively. For IEQ counting, islets or EEOs were enriched in complete RPMI medium at a density of over one islet or EEO per μL. After resuspension, images of five 20-µL droplets were captured and analyzed in ImageJ. Image files were converted to 8-bit format and binarized, followed by threshold adjustment to acquire processed images of the islets. Particle analysis was then applied to extract all the area information of the islets. Based on the equation for the area of a circle, an effective diameter can be calculated for an islet or EEO, and the IEQ was calculated based on the IEQ factor (Table S3). For the number fold change and IEQ yield calculation of islet downsizing, the number (N_d0_) and IEQ (IEQ_d0_) of islets were counted right after islet isolation. For the native islets, after overnight recovery, islets with necrotic cores were removed, and the number (N_d1_) and IEQ (IEQ_d1_) of the remaining islets was calculated on day 1. The fold change of islet number equals to N_d1_/ N_d0,_ and the yield is calculated by IEQ_d1_/ IEQ_d0_ (%). For the EEOs, all the EEOs were quantified on day 5, and the calculation of number fold change (N_d5_/ N_d0_) and yield (IEQ_d5_/IEQ_d0_ (%)) was the same as that of native islets. After 5 days’ recovery, EEOs were further purified prior to in vivo transplantation study. EEO-40 and EEO-70 islets were obtained following the step of filtering through 20 μm cell strainers to remove any remaining cell debris. The size distribution of EEO-40 and EEO-70, and their number and IEQ fraction against size were characterized. The immunofluorescence imaging, intracellular flow cytometry (86) and viability under hypoxic conditions are in Supplementary methods (biological n=3).

### Encapsulation of islets or EEOs

Islets or EEOs were precipitated in a 500 μL EP tube followed by supernatant removal. Islets or EEOs were mixed with 2% alginate solution in 0.9% saline at a density of 15 IEQ/μL. 20 μL of alginate suspension with islets or EEOs at a dose of 300 IEQ was added into a PDMS mold (4 mm × 10 mm x 0.5 mm) with a PET mesh (3 mm x 9 mm x ∼0.1 mm) inside. Then, the mold was transferred into the crosslinking buffer containing 95 mM CaCl_2_ and 5 mM BaCl_2_ for 8 min. The device was subsequently removed out of the mold and incubated in the crosslinking buffer for 2 min. Then, the device was soaked in 1 mL of saline solution 3 times for 1 minute each time. Finally, the device was put into the FBS-free RPMI medium with 1% P/S for subcutaneous transplantation in diabetic mice (Supplementary methods).

### Intraperitoneal glucose tolerance test (IPGTT)

At week 5, mice were fasted overnight and intraperitoneally injected with the glucose solution at a dose of 2 g/kg. The BG at 0, 15, 30, 60, 90 and 120 min were recorded, and the area under the BG-time curve was analyzed.

### Device retrieval and characterization

After device retrieval, each device was cut into two pieces. One half with the intact tissue capsule was directly fixed by 4% FPA for histology study. The other half piece was processed by removing the connective tissue surrounding the alginate device, which was followed by ex vivo GSIS in normal oxygen environments. Before testing, the device was washed by KBR buffer three times and transferred to 24-well plate, then incubated for 2 hours at 37 °C, 21% O_2_ and 5% CO_2_, and then in low glucose concentration (2.8 mM) for 1.5h, followed by high glucose concentration (16.7 mM) for 1.5h at same conditions. The solution was stored in the −80 °C until conducting ELISA (ALPCO, 80-INSRT-E01) as the manual instruction. After GSIS, devices pieces were stained with live/dead staining kit. The black-and-white images were analyzed for the ratio of healthy islets or EEOs indicated by normal morphology and uniform light color without a dark center. Finally, the pieces were fixed in 10% Formalin PBS for 24 hours and placed in 70% ethanol to prepare for histology analysis (Supplementary methods). Representative samples were stained by dithizone (DTZ). DTZ stock solution was prepared at 10 mg/mL in dimethyl sulfoxide. The working solution was prepared by diluting stock solution with PBS at 1:100 ratio and filtering through a 0.22 μm filter. Devices were washed with PBS three times and stained with DTZ working solution at 37 °C for 30 minutes, followed by another session of PBS washing. Devices were then imaged by a stereomicroscope.

### Oxygen-transport Modeling

Mass-transfer models were developed for studying oxygen tension distributions at a steady state in a single endocrine object as a whole and endocrine objects (islets or EEOs) encapsulated in the alginate devices based on similar methods and assumptions (34). Two models were implemented in COMSOL Multiphysics®: one with a single endocrine object as the whole domain, and the other with endocrine objects in a flat alginate device (parameters and boundary conditions are provided in the supplementary methods and Table S2).

### Statistical Analysis

The statistical analysis was conducted using GraphPad Prism (version: 8.0.1). The quantitative data was expressed as Mean ± Standard deviation (SD). Intergroup comparisons were assessed using unpaired Student’s t-test, One-way ANOVA test, or Two-way ANOVA test. Statistical significance was defined as P < 0.05. *P<0.05, **P <0.01, *** P < 0.001, ****P<0.0001. Detailed statistical analysis was summarized in Table S7.

## Acknowledgments

The authors thank Cornell Histology Core Facility for histological cross-sectioning and staining; Cornell Flow Cytometry Core Facility for the technical support; Cornell BRC Imaging Facility (NIH S10OD025049) for laser confocal data. Extended thanks to Shitong Zhao at Cornell University for the material support during early investigation. This work was supported by the National Institute of Diabetes and Digestive and Kidney Diseases (RO1DK143077) and Gates Foundation (INV-062260). Some schematics were created with BioRender.com.

## Data and materials availability

All data associated with this study are presented in the paper and/or the Supplementary Materials.

## Author Contributions

Conceptualization: K.W. and M.M. Methodology: K.W., K.O., M.J., and J.D. Investigation: K.W., K.O., M.J, M.L., J.D., and B.W. Visualization: K.W, K.O., M.J., and M.L. Supervision: K.W., K.O., M.J., M.L., and M.M. Writing—original draft: K.W., M.L. Writing—review and editing: K.W., M.L., M.J., K.O., J.D., B.W., J.A.F., P.T. and M.M. These authors contributed equally: Kecheng Wang, Kento Okada.

## Competing Interest Statement

M.M. is a cofounder of AvantGuard Inc. and Persista Bio. M.M was on leave from Cornell and worked for Vertex Pharma. This work was completed at Cornell, led by K.W. and unrelated to any of the external entities.

