## Supplemental information for "Engineered endocrine objects tolerate hypoxia in macroencapsulation devices and enable subcutaneous engraftment in diabetic mice"

**Contents:**

Supplementary Methods
Figures S1 to S12
Tables S1 to S9

**Supplementary Methods**

**Chemicals.** Calcium chloride dihydrate, barium chloride dihydrate, streptozotocin, citrate acid, Tween 20, Triton-X100, Dithizone and Histopaque®-1077 were purchased from Sigma-Aldrich, sodium alginate (Pronova, SLG100) was purchased from NovaMatrix, Silicone elastomer (SYLGARDTM 184) was purchased from Dow Silicones Corporation. D-glucose anhydrous was purchased from Amresco, 50% Dextrose Injection (USP) was purchased from Hospira, 0.9% Sodium Chloride Irrigation (USP) was purchased from B. Braun Medical. PET mesh (PETKEM 2002) was purchased from SurgicalMesh.

**Islet isolation and purification**

Male or female Sprague Dawley (SD) rats with body weights between 250 g to 350 g were used for islet isolation. Rats were euthanized under anesthesia via exsanguination by cutting the diaphragm and aorta. The pancreas was harvested and perfused with 11 mL of Liberase solution (0.15 mg/mL, in RPMI medium, Roche, 5401020001) and stored on ice. The harvested pancreas was digested within 45 min after perfusion in 37 °C for 28 min. In the middle of the digestion, the tube with the pancreas was shaken and tapped to partially destroy the pancreas. At the end of the digestion, termination buffer (HBSS with 2.5% FBS and 1% penicillin/streptomycin) was added to a total volume of 35 mL. The tube was then shaken and swirled to disperse the pancreas into small nodes. The tube was additionally shaken left to right quickly for 10 times in 10 seconds until fine tissue particles appeared, followed by another 10 times of strong shaking from left to right to release the islets. Following this, islets were washed twice with termination buffer and centrifuged at 230G for 2 min. The cell pellet was dispersed and filtered through a 425 μm sieve, and additional termination buffer was added to increase the volume to 40 mL. The cells were collected after an additional centrifugation at 230G for 2 min. Afterwards, the supernatant was decanted and 20 mL of Histopaque-1077 was added to the cell pellet. The cell pellet was thoroughly shaken left to right about 10 times to redisperse the pellet, and 10 mL of termination buffer was gently loaded on the top. The cell pellet was centrifuged at 900G for 20 min at 1 acceleration and 0 deceleration in 4 °C. All the supernatant from this centrifugal session was filtered by a 70 μm cell strainer, where islets remained on the mesh and were rinsed with 1 mL of termination buffer 5 times to remove residual Histopaque. Islets were then transferred into the termination buffer in a 10 cm petri dish on ice for islet pick-up under a bright-field microscope. The filtrate would be further filtered by a 40 μm cell strainer and all the islets between 40 μm and 70 μm were also collected. All the islets were manually picked twice to increase the purity of the islets. After this step, the islets were ready for the downsizing process or put directly into the incubator for recovery overnight and transplantation. For in vivo experiments, only male SD rat islets were used as recipients for consistency. To maximize islet yield for transplantation, if there were any tiny islets or islet fragments observed which cannot be separated from the exocrine debris manually, the exocrine debris along with these tiny islets were all put into complete RPMI culture medium and incubated overnight. These tiny islets would then be purified by gradient density centrifugation, using the same procedure as for downsized islets. These tiny islets and islet fragments were usually less than 50 μm and were later combined with the majority of islets for in vivo transplantation.

For mouse islet isolation, 2 mL of Liberase (0.15 mg/mL) was perfused into the BALB/c mice and the pancreases were digested in 37 °C water bath for 18 min, with a short shaking by hand occurring at the 9 min. The other procedures were the same as the rat islet isolation.

**Preparation of pseudo-islets and pipetting-disrupted islet clusters.** Native rat islets after one day of recovery were chosen for pseudo-islet preparation. 600 IEQ of islets were collected and washed once with PBS. Then, 0.5 mL of 0.125% Trypsin was applied to the islet pellets. Islets were dispersed well and incubated in a 37 °C water bath for 10 min, and pipetted 10 times every 2 min via 200 μL tips. Afterwards, 1 mL of complete medium was added to stop islet dissociation. Islet cells were collected by centrifugation at 500G for 3 min followed by two repeats. Finally, 1 mL of complete medium was added for cell counting. Collected islet cells were seeded into one well (1200 microwells, low number) or three wells (3600 microwells, high number) in the 24-well AggreWell™ plate. Cells were cultured for 5 days, and the culture medium was half changed every 2 days. Pseudo-islets were collected and counted on day 5. To prepare pipetting-disrupted islet clusters, native rat islets were selected and partially digested by 0.0625% trypsin at room temperature for 5 min on day 0, and then were pipetted by 200 μL tips until less than 20% of the islets were intact (61).

**Immunofluorescence staining of islets and EEOs**. Islets or EEOs were washed with PBS 3 times in an EP tube and then 1 mL of 4% paraformaldehyde (PFA) was added for 30 min at room temperature (RT). After fixation, islets were washed with PBS for 3 times. Then, the islets were dispersed into 1 mL of 0.5% Triton X-100 PBS at RT for 25 min. After PBS washing, islets were incubated in a block buffer (0.1% Tween 20, 10% donkey serum and 1% BSA in PBS) at RT for 1h. Each sample was divided into two parts, one for insulin and glucagon staining, and the other for somatostatin staining. 200 μL of anti-insulin antibody (Abcam, ab63820) and anti-glucagon antibody (Invitrogen, 14-9743-82), or anti-somatostatin antibody (Invitrogen, 14-9751-80) with 200x dilution was incubated in the block buffer at 4 °C overnight. Islets were washed with 1 mL of PBST (0.1% Tween 20 PBS) for 3 times (5 min per time) at 150 rpm. The secondary antibodies were centrifuged at 3000 rpm for 30 seconds before use. Secondary antibodies, Donkey anti-Mouse 488 (Invitrogen, A-21202) and Donkey anti-Rabbit 568 (Invitrogen, A10042) were diluted in 1% BSA solution at the ratio of 1:400 and added into the EP tubes. Islets were incubated for 1 hour at RT in the dark, then washed by PBST to remove uncombined second antibody. DAPI was added for nuclear staining. Finally, islets were imaged using a confocal laser scanning microscope (Zeiss LSM880).

**Intracellular protein staining for flow cytometry.** The intracellular staining protocol was adapted from a previously reported method (86) with minor modifications. Islets were washed with PBS once and digested into single cells in 1 mL of 0.25% trypsin at 37 °C. Native islets were digested for 45 min with gentle pipetting every 15 min until the islets were dispersed into single cells. For the EEOs, due to their smaller size, 25 min was enough for dissociation, and pipetting occurred at the 15 min and the 25 min. Dissociation was terminated by adding complete medium, and cells were collected by centrifugation at 300 G for 3 min at room temperature. Islet cells were washed with PBS and stained with fixable violet live/dead staining dye (Invitrogen, L34963) on ice for 30 min. The supernatant was removed and the cells were washed with PBS once after this. Then, 4% PFA was added to fix the cells on ice for another 30 min. Fixed cells were permeabilized and blocked for 45 min at 4 °C in PBS containing 0.1% (vol/vol) Triton X-100 and 5% (vol/vol) donkey serum. Each cell sample was divided into three aliquots, each of which was stained separately for insulin, glucagon, or somatostatin. Cells were incubated overnight at 4 °C with primary antibodies against insulin (Abcam, ab63820), glucagon (Invitrogen, 14-9743-82), or somatostatin (Invitrogen, 14-9751-80) at 1:300 dilution. After washing, cells were incubated on ice for 2 h with the appropriate secondary antibodies, donkey anti-mouse Alexa Fluor 488 (Invitrogen, A-21202) or donkey anti-rabbit Alexa Fluor 568 (Invitrogen, A10042). Cells were subsequently washed, filtered through flow-cytometry tube strainers, and analyzed by flow cytometry (biological n=3). Unstained, single-stained, and secondary-antibody-only controls were included.

**Static glucose stimulated insulin secretion (GSIS) in vitro.** Glucose was dissolved to a concentration of 2.8 mM as low-glucose condition and to 16.7 mM as a high-glucose condition in Krebs-Ringer Bicarbonate (KRB) buffer (98.5 mM NaCl, 4.9 mM KCl, 2.6 mM $CaCl_2.H_2O$, 1.2 mM $MgSO_4.7H_2O$, 1.2 mM $KH_2PO_4$, 25.9 mM $NaHCO_3$, 0.10% BSA, and 20 mM HEPES (Gibco)). Native islets (IS-N), EEOs or pipetting-disrupted islet clusters (IS-P) (~50 IEQ) were placed in 3 µm Transwell inserts (Corning, 353096) and rinsed by 0.5 mL KRB buffer twice (86). After the KRB buffer was drained by wicking, IS-N, EEOs or IS-P in the inserts were put into a well of 1 mL of 2.8 mM glucose buffer for 1 hour at 37 °C in normal conditions of 21% $O_2$ and 5% $CO_2$, or in hypoxic conditions of 1% $O_2$ and 5% $CO_2$. Following this, they were put into a second well of 2.8 mM glucose solution for another hour. Then, the islets were transferred to a well with 16.7 mM glucose solution for 1 hour. The second 2.8 mM glucose solution and the 16.7 mM glucose solution were stored in -80 °C for insulin analysis. ~90 IEQ of IS-N, EEO-70-r and EEO-40-r islets were digested into single cells and the cell number per IEQ was counted (Table S8). The insulin content was measured by ELISA (ALPCO, 80-INSRT-E01) and normalized by cell numbers. The insulin secretion index refers to the ratio of insulin released under high-glucose stimulation compared to that released under low-glucose conditions.

**Viability study of islets and EEOs under hypoxic conditions.** IS-N, EEO-70, and EEO-40 islets were put into 35 mm petri dishes with 2 mL complete RPMI medium for 24 hours at 1% oxygen level and normal conditions. All the islets or EEOs were picked out for live/dead staining with 1000x dilution of 1 mM Calcein AM Viability Dye (Invitrogen, 65-0853-78) and 20x dilution of 1 mg/mL Propidium (BIOTIUM, 40017) in PBS for 30 min in a 37 °C incubator and then used for imaging. The green and red stained areas of the islet were quantified by ImageJ, and the live area percentage was calculated using green area/(green area + red area). The images were first split into channels that contained only red or green color. We standardized the image analysis process by using the thresholding method “moments” which produced the most accurate detection of coloring compared to other methods for all images, and the total area of green/red was analyzed for all images. For the live/dead staining of the islets in the retrieved alginate devices, they were rinsed by PBS 3 times and stained in 24-well plates for 30 min in a 37 °C incubator before imaging.

**Subcutaneous transplantation of islets or EEOs.** C57BL/6J mice received streptozotocin (STZ) intraperitoneally at a dose of 150 mg/kg. Mice with blood glucose measurements above 300 mg/dL for at least three consecutive days were considered diabetic. The islet or EEO devices were transplanted into the subcutaneous space at the upper right dorsal back area of the mice with an incision of 1 cm made in the skin for transplantation. The incisions were closed by 5-0 sutures.

**Histology study.** The tissue was embedded, sectioned, and stained using H&E (hematoxylin and eosin) by the Cornell core histology facility. Antigen retrieval in citric acid buffer (10 mM citric acid and 0.05% Tween 20, pH 6.0) was conducted at 90 °C for 20 min. Then, the slides were cooled for 10 min at RT, washed in $H_2O$ twice, and in PBS twice. The slides were pre-blocked with 5% donkey serum in PBS for 60 min at RT. Primary antibodies: anti-insulin antibody (Abcam, ab63820), anti-glucagon antibody (Invitrogen, 14-9743-82), and anti-somatostatin antibody (Invitrogen, 14-9751-80) were diluted (1:200) for islet staining; anti-CD31 (Invitrogen, MA5-37858) and anti-a-SMA (Invitrogen, 14-9760-82) were diluted in DB (1:200) for blood vessel staining, and cultured at 4 °C overnight. Secondary antibodies include Donkey anti-Mouse 488 (Invitrogen, A-21202) and Donkey anti-Rabbit 568 (Invitrogen, A10042) with the dilution of 1:400. Before DAPI staining, Vector® TrueVIEW was added and incubated for 5 min to remove autofluorescence.

**Blood glucose measurement.** For the first week after islet graft transplantation, the BG (blood glucose) was measured daily by glucose monitor (Ascensia CONTOUR NEXT EZ). Graft failure was defined as BG above 300 mg/dL for two consecutive days post one-week transplantation, whereas functional engraftment was indicated by BG below 200 mg/dL for two consecutive days without subsequent graft failure during the observation period. If a mouse is able to maintain BG below 200 mg/dL for at least two consecutive days, the BG was measured two or three times a week. After device retrieval, the BG of the engrafted mice was monitored for another 2 weeks to confirm the function of the implants. When mice exhibited severe morbidity following device retrieval, they were euthanized immediately following the animal protocol.

**Oxygen-transport Modeling.** Mass-transfer models were developed for studying oxygen tension distribution at steady state in a single endocrine object as the whole domain and endocrine objects encapsulated in the alginate devices based on reported methods and assumptions (34). Oxygen has a poor solubility in aqueous media such as in body tissue, culture medium and alginate hydrogel. To be simple and convenient, for this study it is assumed that there was no oxygen solubility difference among endocrine objects and alginate hydrogel in the simulation model. The conservation equation for a diluted species is given by:

$$\frac{\partial c}{\partial t} = D_{O_{2,i}} \nabla^2 c - V_{O_{2,i}} \qquad (1)$$

where $c$ is the dissolved oxygen concentration, t is the time, $D_{O_{2,i}}$ is the oxygen diffusion coefficient in subdomain $i$ , and $V_{O_{2,i}}$ is the volumetric oxygen consumption rate of endocrine objects in subdomain $i$. In steady state, the oxygen concentration will not change over time, as shown in the steady state equation:

$$\frac{\partial c}{\partial t} = 0 \qquad (2)$$

Henry's law allows us to express the dissolved oxygen concentration in terms of the partial pressure $p$ as shown below:

$$c = \alpha_{O_{2,i}} p \qquad (3)$$

where $\alpha_{O_{2,i}}$ is the partial pressure-dependent solubility coefficient of oxygen in the subdomain $i$. The oxygen consumption rate can be governed by Michaelis-Menten kinetics and a step-down function based on the lack of oxygen consumption in the necrotic area, as shown by:

$$V_{O_{2,i}} = \begin{cases} 0, & p < p_N \\ \frac{pV_{max}}{p+K_v}, & p \geq p_N \end{cases} \qquad (4)$$

where $V_{max}$ is the maximum basal OCR, $K_v$ is the half-maximal coefficient and $p_N$ is the threshold oxygen tension of cell survival. There is no oxygen consumption in the domain of culture medium and alginate hydrogel.

Two models have been implemented in COMSOL Multiphysics®: one with a single endocrine object as the whole domain, and the other with endocrine objects in a flat alginate device. To explore the minimal peripheral oxygen tension that could support endocrine objects without necrosis development, single endocrine object with different diameters and different peripheral boundary oxygen tensions ($p_{ext}$) were used for computation to obtain the minimal peripheral oxygen tension-endocrine object diameter curve (diameter: 1 μm resolution). For the endocrine objects in the alginate device for in vivo simulation, the model is based on the beforementioned "SHARP" model, with alterations in the endocrine object distribution parameters that were fitted by the Weibull distribution, and boundary oxygen tension ($p_{ext}$) was set as 1% $O_2$. The parameters were listed in Table S2 and Table S9.

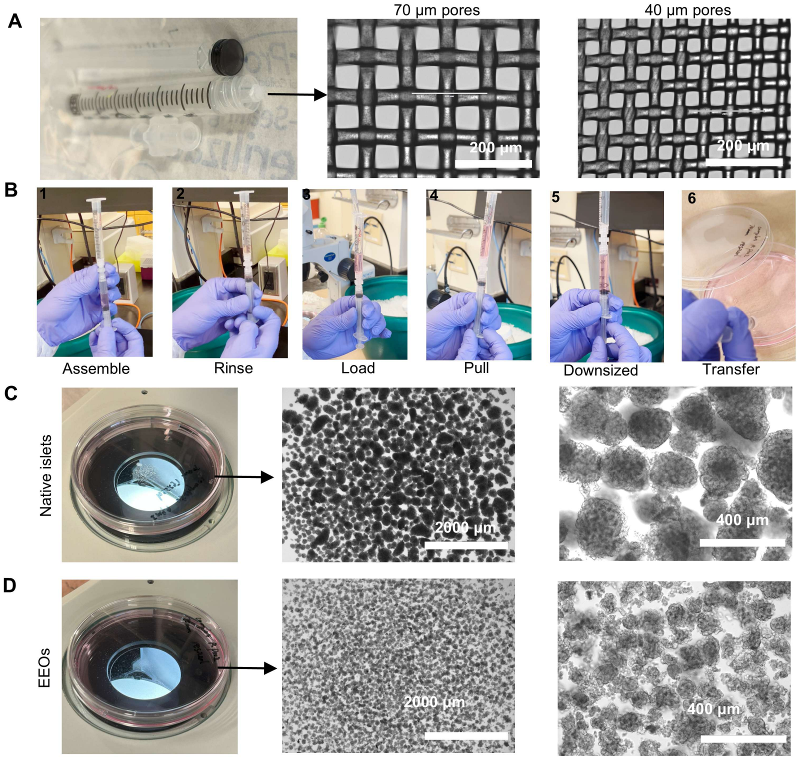


**Fig. S1: Islet downsizer enables efficient preparation of rat EEOs.** (A) The digital image of the islet-downsizer with a nylon mesh. (B) Illustration of the islet downsizing procedures. Digital images of native rat islets before (C) and after (D) downsizing.

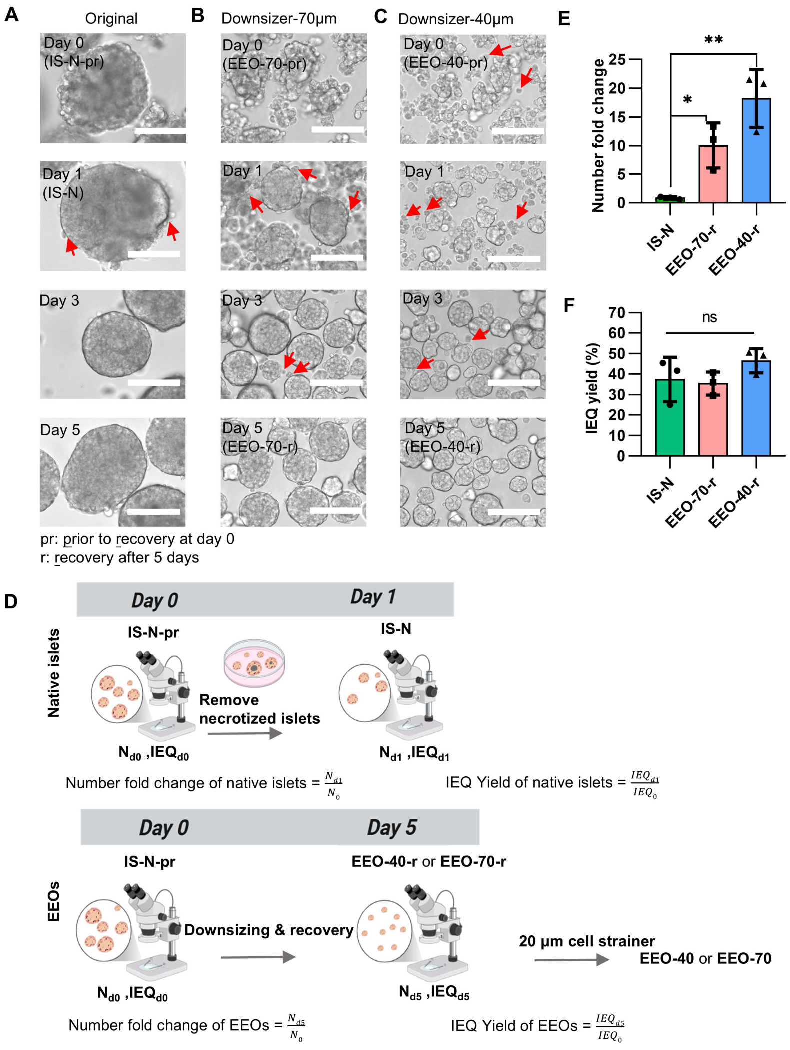


**Fig. S2: Spontaneous recovery and quantification of native and EEOs over time.** Morphology of rat islets over time: native islets (A), EEOs prepared by downsizer-70 µm (B) and EEOs prepared by downsizer-40 µm (C). The red arrows point to dead cells or cell debris. Scale bar: 100 µm. (D) Schematics of quantification of native islets and EEOs in number fold change and IEQ yield. (E) The fold change of islet or EEO numbers. (F) The IEQ yield. Data is presented as mean ± SD, analyzed by one-way ANOVA. ns, not significant; $^*P < 0.05$; $^{**}P < 0.01$.

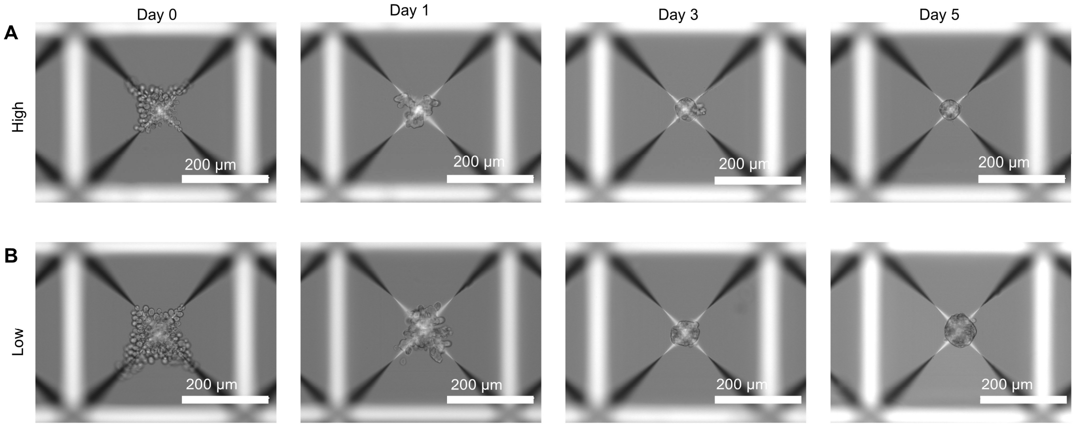


**Fig. S3: Pseudo-islets preparation by single islet cell reaggregation in low (1200) and high (3600) number of microwells.** Representative black-and-white images of pseudo-islets formation over time in high number of microwells (A) and low number of microwells (B).

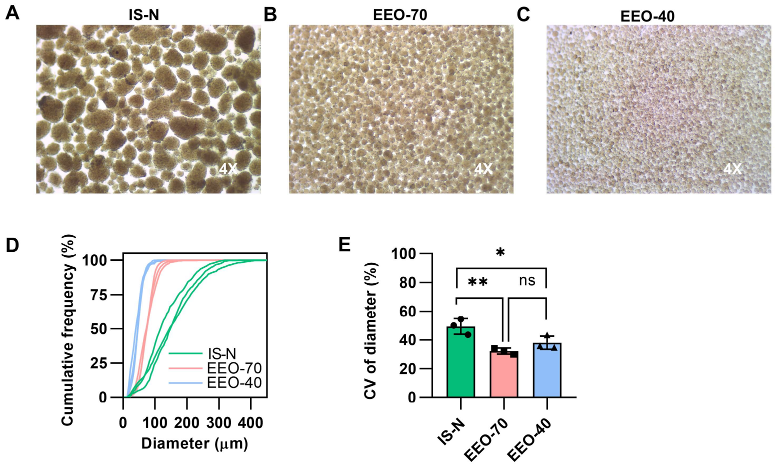


**Fig. S4: Downsized rat islet clusters recovered to a round and smooth morphology with a narrow size distribution after recovery culture.** Stereoscopic images of (A) native islets (IS-N), (B) EEO-70, and (C) EEO-40 after 5 days recovery culture and filtered by 20 µm cell strainers. (D) Size cumulative frequency of IS-N, EEO-70 and EEO-40 (Biological n=3, islet number > 290 in each group). (E) inter-batch coefficient of variation (CV) of diameters. Data is presented as mean ± SD, analyzed by one-way ANOVA. ns, not significant; *P < 0.05; **P < 0.01.

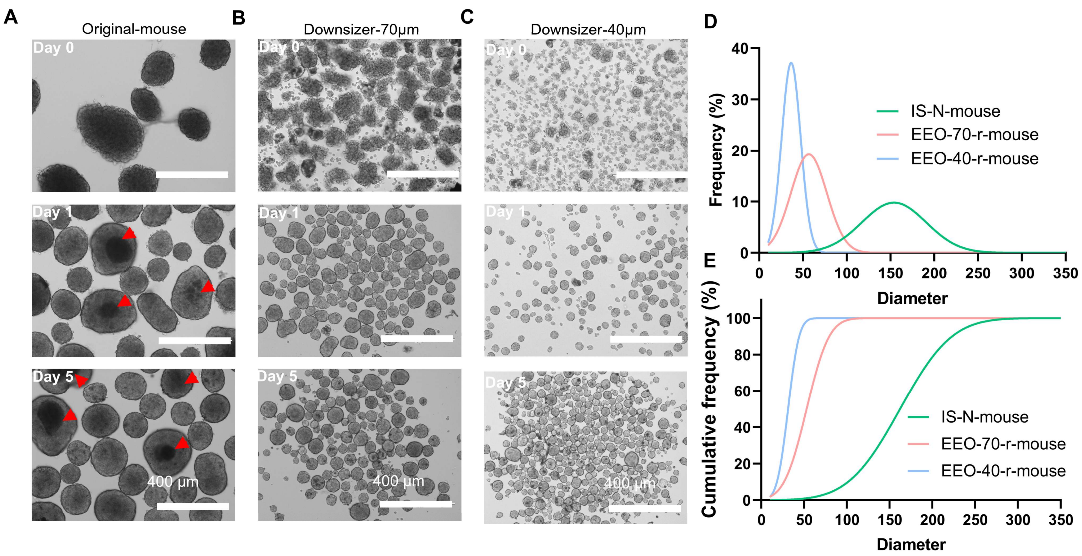


**Fig. S5: Islet downsizing strategy extends to mouse islets.** Islets isolated from BALB/c mice underwent the same downsizing process as rat islets. The morphology changes of (A) mouse native islets, mouse EEOs by 70 µm (B) and 40 µm pores (C) at day 0, day 1 and day 5. Red arrows indicate islet necrotic cores. (D) Size distribution and cumulative frequency (E) of IS-N-mouse, EEO-70-r-mouse and EEO-40-r-mouse.

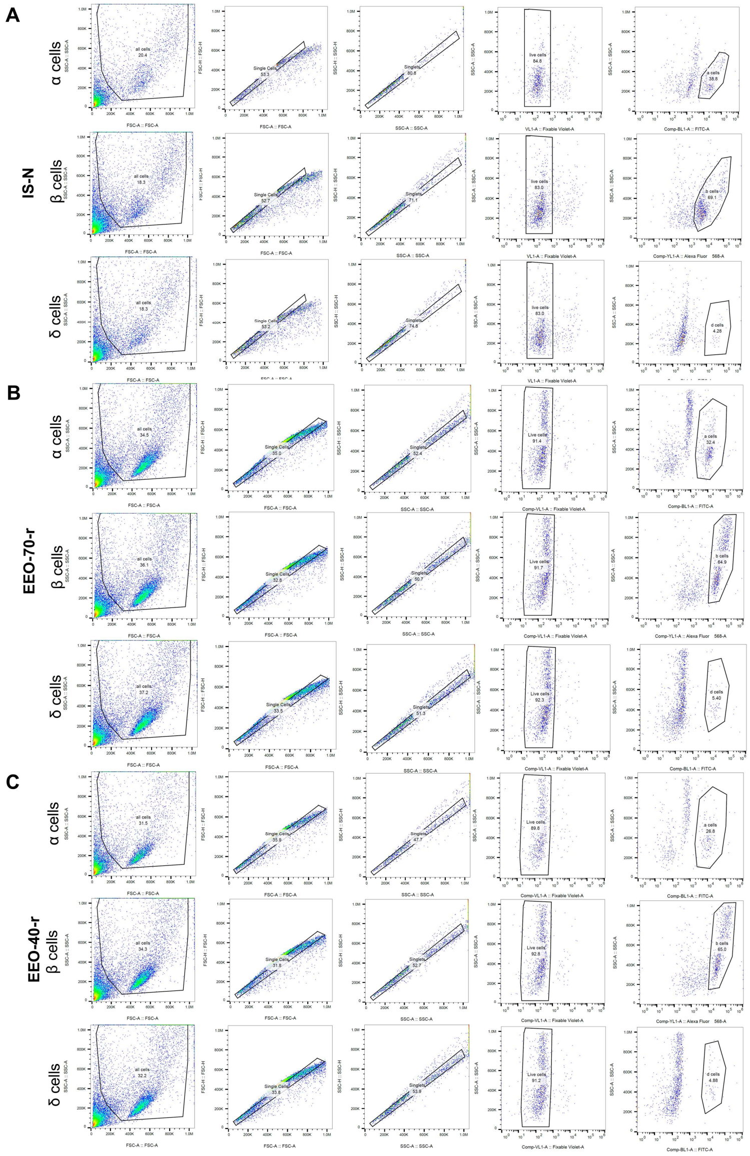


**Fig. S6: Flow cytometry data of α, β, δ cells.** Debris and doublets were removed for analysis in native islet cells (A), EEO-70-r (B) and EEO-40-r (C) (biological n=3).

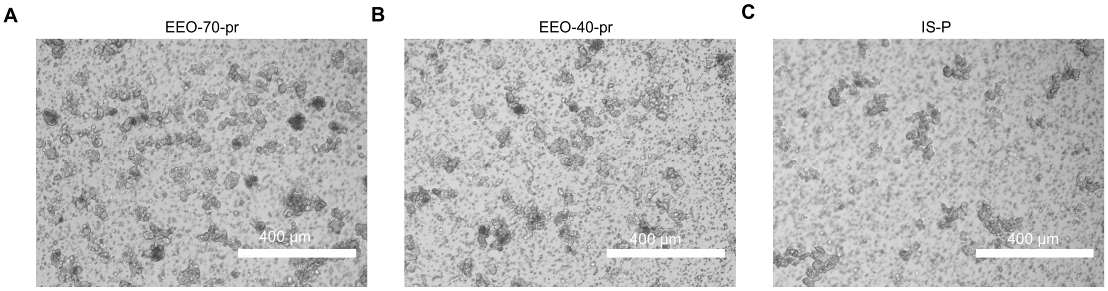


**Fig. S7: Representative black-and-white images of EEO-70-pr, EEO-40-pr and IS-P (pipetting-disrupted islet clusters on day 0) on the Transwell membrane for GSIS.**

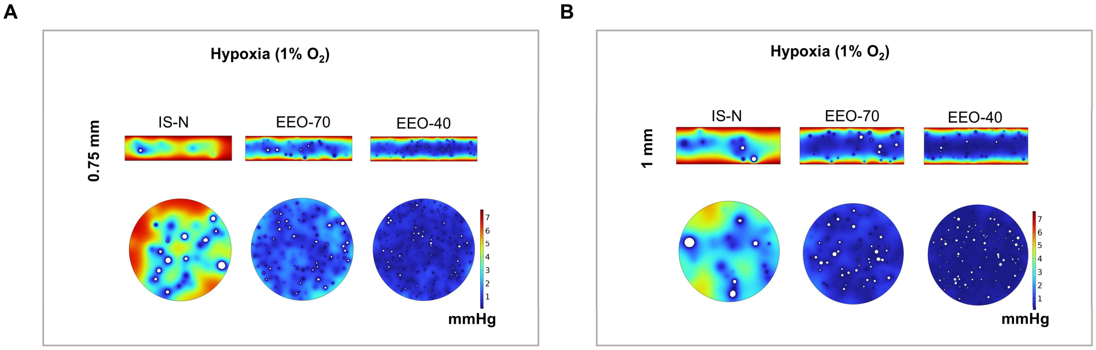


**Fig. S8: The effect of encapsulation device thickness on oxygen tension distribution simulated by COMSOL.** The oxygen tension distribution of IS-N, EEO-70 and EEO-40 in central vertical (top) and horizontal (bottom) cross-section of the flat alginate device with a peri $O_2$ tension of 7.6 mmHg (1% $O_2$) at a thickness of 0.75 mm (A) and 1 mm (B) in an islet loading density of 15,000 IEQ/mL. The white area indicates the necrotic area ($O_2$ tension < 0.08 mmHg).

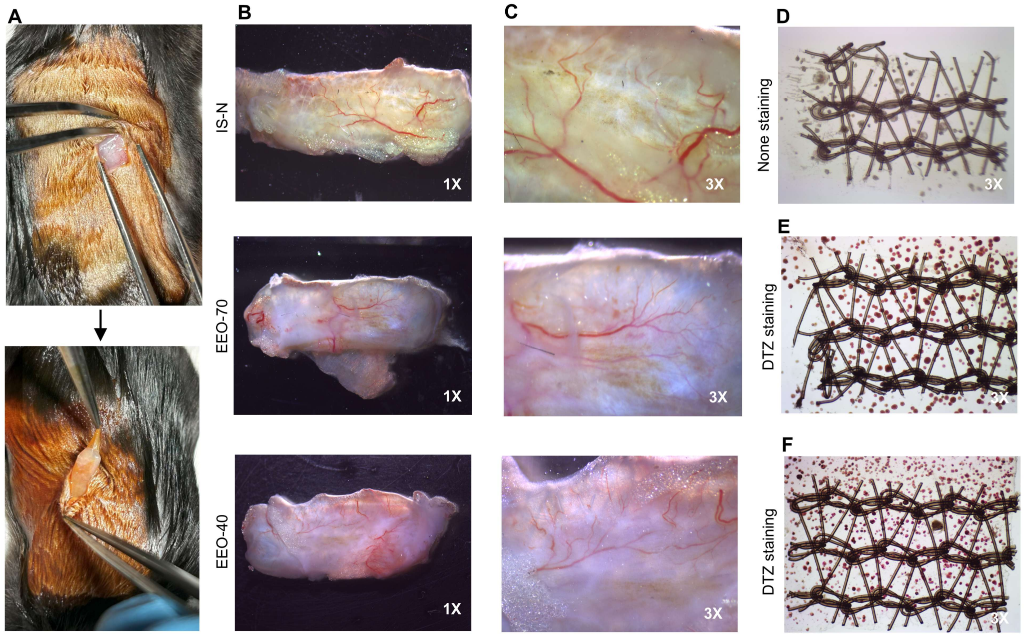


**Fig. S9: Alginate device maintained intact at retrieval and the EEOs within it were DTZ-positive.** (A) Image of the retrieval procedure. Digital images of the tissue capsule after device retrieval and blood vessel under 1x (B) and 3x (C) magnification. (D) Stereomicroscopic image of one unstained native islet device peeled from the capsule. DTZ staining of the retrieved EEO-70 (E) and EEO-40 (F) device after 5-week subcutaneous transplantation.

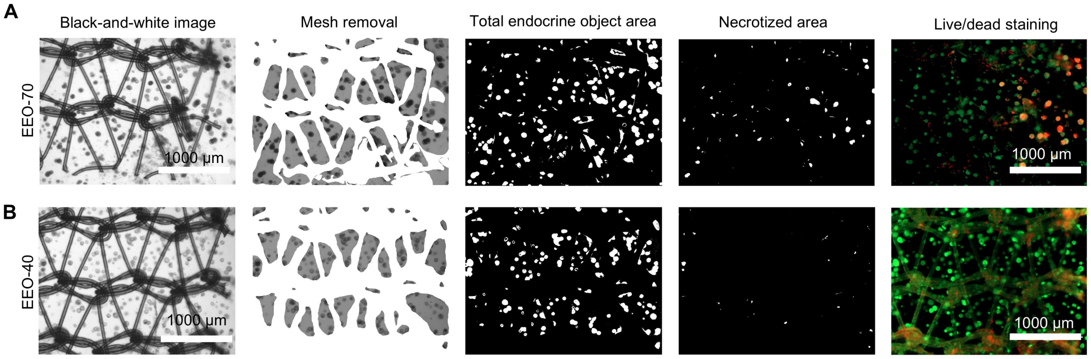


**Fig. S10: Necrotic area percentage calculation by black-and-white images and validation by related live/dead staining images.** Necrotic areas of EEO-70 (A) and EEO-40 (B) in the retrieved flat alginate devices were quantified based on black-and-white images after the mesh removal. The total endocrine object area and dark necrotic centers were captured by ImageJ.

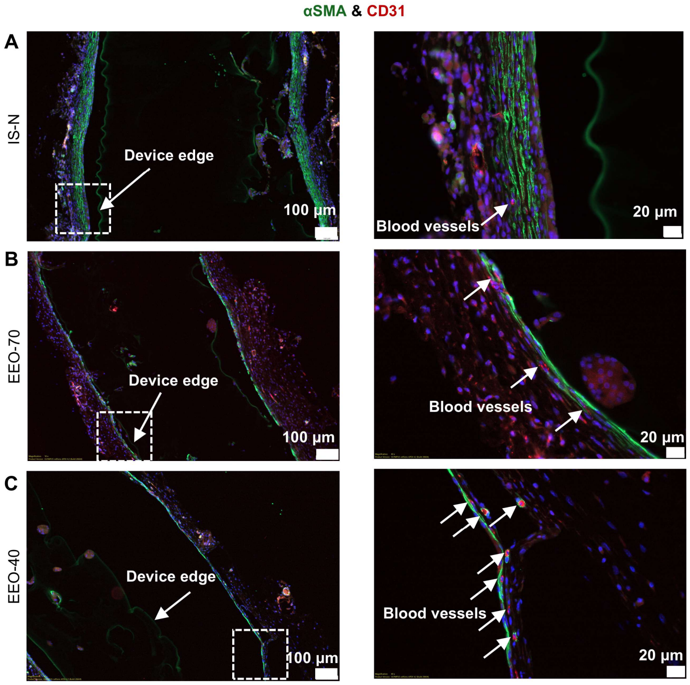


**Fig. S11: Vascularization at the tissue-device interface after subcutaneous transplantation.** Representative immunofluorescence images of αSMA and CD31 staining in tissue surrounding the retrieved alginate devices containing IS-N (A), EEO-70 (B) and EEO-40 (C).

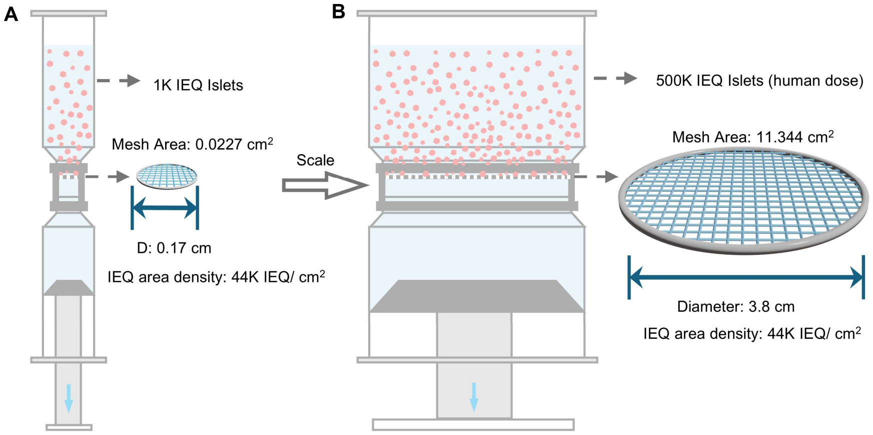


**Fig. S12: Conceptual scale-up of the islet downsizer for future human islet processing.** (A) The illustration of the laboratory-scale islet downsizer used in the present study. (B) The scalability of islet downsizer. Conceptual scale-up of the islet downsizer for processing a clinically relevant quantity of islets. Assuming the same islet loading per unit mesh area, processing 500K IEQ would require a mesh area of approximately 11.344 cm², corresponding to a circular mesh with a diameter of approximately 3.8 cm.

### Table S1. Subcutaneous oxygen tension in different species.

| Animal | Subcutaneous sublocation | Measurement method | $pO_2$ (mmHg) | References |
|---|---|---|---|---|
| Human | superficial region of the skin | Clark type oxygen probe | ~8.0 mmHg | 63 |
| Human | dermal papillae | Clark type oxygen probe | ~24.0 mmHg | 63 |
| Human | above the subpapillary plexus | Clark type oxygen probe | ~35.2 mmHg | 63 |
| Non-human primate | N/A | Clark type oxygen probe | ~39.0 mmHg | 64 |
| Rat | dorsal column | Electronic paramagnetic resonance (EPR) oximetry | 22.8-38 mmHg | 65 |
| Rat | paravertebral space | EPR | ~40 mmHg | 66 |
| Rat | N/A | flexible wire-type $O_2$ probe | 48.5 mmHg | 67 |
| Rat | dorsal | $^{19}$F-MRS oximetry | ~10 mmHg | 68 |
| Rat | N/A | optical oxygen sensors | ~4.1 mmHg | 69 |
| Mouse | N/A | EPR | 1.41-26 mmHg | 70 |
| Mouse | Abdomen | EPR | 1.68-21.23 mmHg | 32 |

### Table S2. Simulation model parameters.

| Symbol | Description | Value (unit) |
|---|---|---|
| $\alpha_{O_{2,i}}$ | Partial pressure-dependent solubility coefficient of oxygen in domain $i$ | $1.24 \times 10^{-3}$ (mol m$^{-3}$ mmHg$^{-1}$) |
| $D_{O_{2,h}}$ | Diffusion coefficient of oxygen in hydrogel | $2.70 \times 10^{-9}$ (m$^2$ s$^{-1}$) |
| $D_{O_{2,c}}$ | Diffusion coefficient of oxygen in islet cells | $2.00 \times 10^{-9}$ (m$^2$ s$^{-1}$) |
| $D_{O_{2,m}}$ | Diffusion coefficient of oxygen in culture medium | $3.00 \times 10^{-9}$ (m$^2$ s$^{-1}$) |
| $V_{max}$ | Maximum oxygen consumption rate (per unit volume) in islets | 0.034 (mol m$^{-3}$ s$^{-1}$) |
| $K_v$ | Half-maximal coefficient of oxygen consumption in islets | 0.80 (mmHg) |
| $p_N$ | Threshold oxygen tension required for cell survival (below which oxygen consumption are ceased) | 0.08 (mmHg) |
| $p_{ext}$ | Boundary $pO_2$* | 0.1-38 (mmHg) |

* A default value of 7.6 mmHg was used for subcutaneous in vivo simulation.

### Table S3. IEQ conversion factor.

| Islet size group (d, μm) | IEQ conversion factor |
|---|---|
| < 50 | 0.028 (34) |
| 51-100 | 0.167 |
| 101-150 | 0.648 |
| 151-200 | 1.685 |
| 201-250 | 3.500 |
| 251-300 | 6.315 |
| 301-350 | 10.352 |
| > 350 | 15.833 |

### Table S4. Yield of islets, EEOs and pseudo-islets (IS-pseudo) (n=3).

| Islet | $IEQ_0$ | $IEQ_{5\ or\ 1(IS\text{-}N)}$ | Yield (%) | Yield coefficient of variation (%) |
|---|---|---|---|---|
| IS-N | 656.59±151.81 | 242.41±89.18 | 37.31±10.81 | 28.98 |
| EEO-70-r | 566.58±108.34 | 201.6±61.22 | 35.33±5.64 | 15.96 |
| EEO-40-r | 545.75±180.55 | 259.35±111.17 | 46.40±5.91 | 12.74 |
| IS-pseudo(L) | 600 | 85.91±59.70 | 14.32±9.95 | 69.50 |
| IS-pseudo(H) | 600 | 189.23±109.86 | 31.54±18.31 | 58.06 |

Yield coefficient of variation (%) = $SD\_{Yield}/Mean\_{Yield}$

### Table S5. Suggested islet treatment dose for different implantation sites.

Subcutaneous space (SQ), intraperitoneal space (IP), Kidney capsule (KC)

| Mouse | Location | Islet donor | Delivery method | Dose (IEQ) | References |
|---|---|---|---|---|---|
| BALB/c | KC | Mouse | Free | 500 | 33 |
| BALB/c | IP | Rat | Encapsulated | 1200 | 19 |
| C57BL/6J | IP | Rat | Encapsulated | 600 | 71 |
| C57BL/6J | IP | Rat | Encapsulated | 500 | 28 |
| C57BL/6J | IP | Rat | Encapsulated | 300-350 | 72 |
| C57BL/6J | IP | Rat | Encapsulated | 300 | 34 |
| SCID Beige | SQ | Human | Free | 3000 | 73 |
| BALB/c nude | SQ | Rat | Encapsulated | 250 | 29 |
| C57BL/6J | SQ, prevascularization | Rat | Encapsulated | 500 | 31 |
| C57BL/6J | SQ, prevascularization | Rat | Encapsulated | 500 | 32 |
| C57BL/6J | SQ, oxygen supply | Rat | Encapsulated | 500 | 23 |
| C57BL/6J | SQ, oxygen supply | Rat | Encapsulated | 850 | 27 |
| C57BL/6 | SQ, antifibrotic drug | Rat | Encapsulated | 500 | 29 |
| C57BL/6J | SQ | Rat | Encapsulated | 300 | this work |

**Table S6. Summary of the sizes of engineered islets for transplantation.**

| Islet source | Method | Size of engineered islets (μm) | Transplant location | Delivery method | Recipient | References |
|---|---|---|---|---|---|---|
| Rat islets | Enzymatic dissociation + spontaneous reaggregation | ~39 | IP | Encapsulated | BALB/c mouse | 44 |
| Rat islets | Enzymatic dissociation + microwell | ~41 | KC | Free | SD rat | 74 |
| Rat islets | Enzymatic dissociation + hanging-drop reaggregation | ~118 | SQ | Free | BALB/c nude mouse | 75 |
| Human islets | Enzymatic dissociation + microwell | ~80-~180 | KC | Free | Rag1$^{-/-}$ mouse | 76 |
| Mouse islets | Fragmented by vigorous shaking or extra collagenase exposure for 10 min | ~128 | KC | Free | C57BL/10 mouse | 36 |
| Rat islets | Fragmented by repeated pipetting after papain exposure for 10 min | ~60 | KC | Free | DA Rat | 61 |
| Rat islets | Fragmented by mesh + recovery culture | ~47 | SQ | Encapsulated | C57BL/6 mouse | This work |

### Table S7. Summary of statistical analysis.

| Figure | Statistical analysis |
|---|---|
| Fig.2G | one-way ANOVA with Tukey's adjusted P valuer for multiple comparisons |
| Fig.2H | one-way ANOVA with Tukey's adjusted P valuer for multiple comparisons |
| Fig.2J | two-way ANOVA, followed by Tukey's multiple comparisons test |
| Fig.2L | two-way ANOVA, followed by Tukey's multiple comparisons test |
| Fig. 3B | Two-way RM ANOVA |
| Fig. 3C | unpaired two-tailed Student's t-test |
| Fig. 3D | one-way ANOVA with Dunnett's adjusted P value for multiple comparisons |
| Fig. 3E | unpaired two-tailed Student's t-test |
| Fig. 3F | one-way ANOVA with Tukey's adjusted P valuer for multiple comparisons |
| Fig. 3I | unpaired two-tailed Student's t-test for the same group with different oxygen conditions |
| Fig. 3I | one-way ANOVA with Tukey's adjusted P value for multiple comparisons in the same oxygen condition |
| Fig. 3J | unpaired two-tailed Student's t-test for the same group with different oxygen conditions |
| Fig. 3J | one-way ANOVA with Tukey's adjusted P value for multiple comparisons in the same oxygen condition |
| Fig. 4B | two-way ANOVA, followed by Tukey's multiple comparisons test |
| Fig. 4H | two-way ANOVA with Greenhouse-Gessier's correction and Tukey's adjusted P value for multiple comparisons |
| Fig. 4I | one-way ANOVA with Tukey's adjusted P value for multiple comparisons. |
| Fig. 4J | unpaired two-tailed Student's t-test |
| Fig. 5F | one-way ANOVA with Tukey's adjusted P value for multiple comparisons |
| Fig. 5G | unpaired two-tailed Student's t-test |
| Fig.S2E | one-way ANOVA with Dunnett's adjusted P value for multiple comparison |
| Fig.S2F | one-way ANOVA with Tukey's adjusted P valuer for multiple comparisons |
| Fig.S4E | one-way ANOVA with Tukey's adjusted P valuer for multiple comparisons |

### Table S8. Islet cell number per IEQ in IS-N, EEO-70-r and EEO-40-r.

| EO | IEQ | EO Number | Cell number | Cell number /IEQ |
|---|---|---|---|---|
| IS-N | 89.83 | 47 | 70500 | 784.8 |
| EEO-70-r | 83.96 | 1410 | 24000 | 285.8 |
| EEO-40-r | 108.12 | 1830 | 36000 | 333.0 |

### Table S9. Weibull distribution parameters of islets for "SHARP" simulation model.

| | Weibull parameters | | |
|---|---|---|---|
| Cell source | $\alpha_n$ | $\beta_n$ | $\overline{d_n}$ (μm) |
| IS-N | 167.5 | 2.14 | 148.8 |
| EEO-70 | 85.9 | 3.6 | 77.6 |
| EEO-40 | 52.8 | 3.06 | 47.4 |
